# Glutamic acid-lysine (EK) rich motif of RabD2 self-associates and regulates pathogenesis through multivesicular bodies pathway in *E. histolytica*

**DOI:** 10.1101/2024.11.21.624624

**Authors:** P Navyaka, Kuldeep Verma

## Abstract

*Entamoeba histolytica*, an enteric pathogen, causes disease by adhering to and destroying the host tissues. The interactions between the parasite and host tissue enable rewiring of the gene expression and global membrane trafficking in the parasite. A fine balance between cargoes/receptors endocytosis and exocytosis is required to establish infection in the host. Multivesicular bodies (MVBs) act as sorting platforms, delivering cargoes/receptors to lysosomes for degradation or secreting their content through plasma membrane fusion. Some of the small GTPases are known to control MVB biogenesis in various organisms. However, the functional contribution of Rab GTPases in MVB biogenesis is poorly studied in *E. histolytica*. Here, we identified a novel atypical protein RabD2, with an N-terminal glutamic acid-lysine rich motif and a C-terminal conserved Rab domain. Our biochemical and cell biological assays provide evidence that RabD2 self-associates, and this interaction is controlled by the N-terminal EK-rich motif and the GTPase activity mutants (in a nucleotide-specific manner). RabD2 localizes on the surface of MVBs and controls their biogenesis. In line with these findings, overexpression of RabD2 upregulates global ubiquitination, directing the down regulation of the heavy chain of GalNAc lectin, ultimately leading to decreased adherence of *E. histolytica* trophozoites to host cells. Thus, amoebic RabD2 is a new class of Rab protein that forms high-ordered self-association variants and regulates the pathogenicity of *E. histolytica* through the biogenesis of MVBs.

**Author Summary:** Amoebiasis is an enteral infection caused by *Entamoeba histolytica* that primarily remains asymptomatic but can eventually result in systemic complications like amoebic dysentery, liver abscess, and pulmonary effusions. Recent studies showed that *Entamoeba* ubiquitin is a robust antigen and linked with the invasive amoebiasis patient samples. Here, we identified a novel RabD2 that self-associates via its glutamic acid-lysine rich motif that causes high ubiquitination levels and biogenesis of multivesicular bodies. We uncovered for the first time that RabD2-mediated ubiquitin-dependent pathway is involved in the down regulation of the notable antigenic marker heavy chain of galactose-N-acetylgalactosamine lectin and thereby controls the parasite adherence to host cells. Further studies on crosstalk between parasite ubiquitination and antigenic receptors downregulation provide insights into how parasites use these strategies to establish the infection in the host intestine.

## Introduction

*Entamoeba histolytica* is a unicellular protozoan parasite which is commonly known to cause intestinal and extraintestinal amoebiasis. The infections are reported across the globe with nearly 100,000 fatalities annually [1]. Moreover, its prevalence is much higher in children below 5 years of age and in low socio-demographic index countries [2]. Endocytosis is one of the fundamental pathways for nutrient uptake, growth, and virulence in this parasite. Multivesicular bodies (MVBs) are a specialized subset of endosomes that are characterized by membrane-bound intraluminal vesicles [3]. Protein localization on MVBs means that the protein is sorted into the MVB lumen or membrane by a process called endosomal sorting.

This process is regulated by various proteins, such as the ESCRT (endosomal sorting complex required for transport) machinery proteins, cytoskeleton members, and vacuolar sorting receptors etc. The genesis of MVBs relies on the sequential recruitment of ESCRT-0, ESCRT-I, ESCRT-II, ESCRT-III and ESCRT-III associated proteins [4]. This process begins with recognition of the mono-ubiquitinated proteins by ESCRT-0 at the cytosolic face of endosomal compartments. The comparative genomic study revealed that ESCRT machinery proteins are highly conserved across the eukaryotic lineage [5]. However, the ESCRT-0 proteins are only found in opisthokonts (metazoans and fungi), which suggests an alternative mechanism for sorting the cargo to MVBs in other organisms, including amoebozoa [6]. The amebic parasite encodes 19 genes of ESCRT machinery proteins, which are devoid of the ESCRT-0 complex proteins [VHS domain (Vps27-Hrs-STAM)], and it is presumed to use an unconventional VHS domain protein/s for the recognition of ubiquitinated proteins [7]. Similar to *A. thaliana* and *D. discoideum*, *E. histolytica* encodes Tom1 and is considered to be a scaffold to interact with ubiquitin and other ESCRT machinery proteins [8, 9]. The ESCRT-I member, EhVps23, an orthologue of yeast Vps23 and mammalian TSG101, is predicted to have a UEV domain (ubiquitin E2 variant domain present in TSG101) which potentially interact with LBPA and ubiquitin [10]. The ESCRT-III protein, EhVps32, interacts with EhVps20 and then recruits EhVps24. EhVps2 has been shown to control the ILV size [11]. Previously, it has been shown that the ESCRT member EhVps23 is involved in vesicular transport, phagocytosis, motility, and secretion. Based on immunoprecipitation and proteomic analysis, it was hypothesized that ESCRT members couple with multiple Rab proteins to take part in MVB functions [10].

More than 100 Rab proteins are encoded by *E. histolytica* [12]. These proteins act as molecular switches and control various membrane transport events such as vesicle tethering, budding, and fusion etc. Rab GTPase cycles between active (GTP-bound) and inactive (GDP-bound) states intrinsically by utilizing the specific regulators such as GEFs and GAPs [13]. Previously, it has been shown that EhRab5 and EhRab7a form unique endocytic pre-phagosomal vacuoles (PPVs) upon interaction with the host cells [14]. These PPVs are indicative of MVB-like structures that transport the acid hydrolases and iron to lysosomes [15].

In higher eukaryotes, Rab7, Rab11, Rab22, and Rab27 are reported to localize on MVBs and control their fusion with lysosomes and plasma membrane [16–19]. Although the direct role of Rab GTPases in MVB biogenesis is poorly studied.

The unbiased transcriptomic screen identified that specifically two sets of Rab GTPases, RabD2 (EHI_164900) and Rab35 (EHI_146510), were upregulated in the virulent strain of *E. histolytica* (HM: 1: IMSS) upon the pathogens interaction with human colon tissue [20]. In a recent microarray-based study conducted in both non-pathogenic (UG10) and pathogenic (HM: 1: IMSS) strains, it was found that RabD2 is upregulated only in the pathogenic strain of *E. histolytica* [21]. Further, upregulation of RabD2 is also linked with cytopathic activity against the host [22]. The RabD2 homologue is absent in non-virulent strains such as *E. dispar* and *E. moshkovskii*. Altogether, it suggests that RabD2 could be an important regulator for the pathophysiology of pathogenic *E. histolytica* strains. Therefore, understanding the function of RabD2 will bring new insights into the pathophysiology of *E. histolytica*. In this study, we have employed quantitative, biochemical and cell biological assays which provide evidence that RabD2 self-associates, and its interaction alone is controlled by the N-terminal glutamic acid-lysine (EK) rich motif and nucleotide status of GTPase. We showed that RabD2 localizes on the MVBs and controls its biogenesis. Furthermore, we found that RabD2 overexpression and activity upregulate global ubiquitination, which leads to the down regulation of the heavy chain of GalNAc lectin (Hgl), a prominent antigenic marker for *E. histolytica* infection [23, 24]. Further, our results showed that adherence of pathogens to host cells is reduced due to the low levels of plasma membrane Hgl. Collectively, our findings demonstrate that the novel RabD2, consisting of the EK-rich motif, undergoes high-ordered association and regulates the *E. histolytica* pathogenicity through MVB biogenesis.

## Results

### Large RabD2 contains an atypical glutamic acid-lysine rich motif and is expressed as SDS-resistant high molecular weight protein bands

To study the details of putative large RabD2 (EHI_164900) from *E. histolytica*, primers were designed based on the hypothetical sequence available on the database (amoebadb.org). The high-fidelity PCR was performed using respective primers (see materials and methods section) and a 795 bp (approximately) amplified product was obtained from the cDNA pool of *E. histolytica* (S1A Fig). The amplified product was cloned in an amoebic expression vector (pEhTex-HA) with an N-terminal 3X hemagglutinin epitope sequence, and it is hypothetically translated into amino acids. The cloned RabD2 hypothetical molecular weight is around 30 kDa, which is slightly higher than the conventional Rab GTPase (20-25 kDa). These high molecular weight Rab GTPases are known as large Rab GTPases [25]. The mammalian Rab44, Rab45, and Rab46 belong to the large Rab GTPases family. Further, Rab2A, together with the known large Rab GTPases Rab44, Rab45, and Rab46, were retrieved from the protein database, and comparative sequence alignment was performed. It was observed that RabD2 consists of a conserved G domain followed by an additional 69 amino acids at the N-terminal (S1B Fig). The N-terminal 69 residues are non-Rab sequences consisting of negatively charged residues (Aspartic acid + Glutamic acid = 31 residues) and positively charged residues (Arginine + Lysine = 19). In addition, this region is rich in glutamic acid (15 residues) and lysine (12 residues) amino acids that are distributed within the N-terminal 69 residues in protein sequences. The specific characteristic abundance of glutamic acid and lysine residues suggested us to name it the EK-rich motif (Fig 1A). Interestingly, bioinformatics analysis also revealed the presence of potential disordered region at the EK-rich motif of RabD2. Further, pEhTex-HA-RabD2 plasmid was transfected, and stable expressing trophozoites were selected in the presence of G418. The pEhTex tetracycline-inducible system allows regulated expression of the target gene/s over two orders of magnitude and is commonly used in expression studies [26–28]. To study the expression of RabD2, transfected trophozoites were incubated with tetracycline, and the lysate was subjected to western blotting using anti-HA antibody and cysteine synthase 1 (CS1) as a loading control (Fig 1B). Surprisingly, our western blot analysis reveals high molecular weight bands including the expected size around 33 kDa (HA tagged with RabD2). Although high molecular weight bands were prominent compared to the expected size. Further, RabD2 expressing trophozoites were incubated without and with tetracycline, and it was observed that the abundance of high molecular weight bands was varying among the experiments, suggesting a dynamic modification under steady-state conditions (Fig 1C). Next we looked into the subcellular localization of HA-RabD2 expressing trophozoites and observed that RabD2 is distributed throughout the cell (Figs 1D and E) and it is mainly localized on the variable-size vesicles (diameter, 0.8 ± 0.12 µm) and vacuoles (diameter, 2.04 ± 0.65 µm). Altogether, our results suggest that RabD2 consist of EK-rich motif, migrates at high molecular weight species, and it is localized on variable sizes of vesicular and vacuolar compartments.

**Fig 1.**
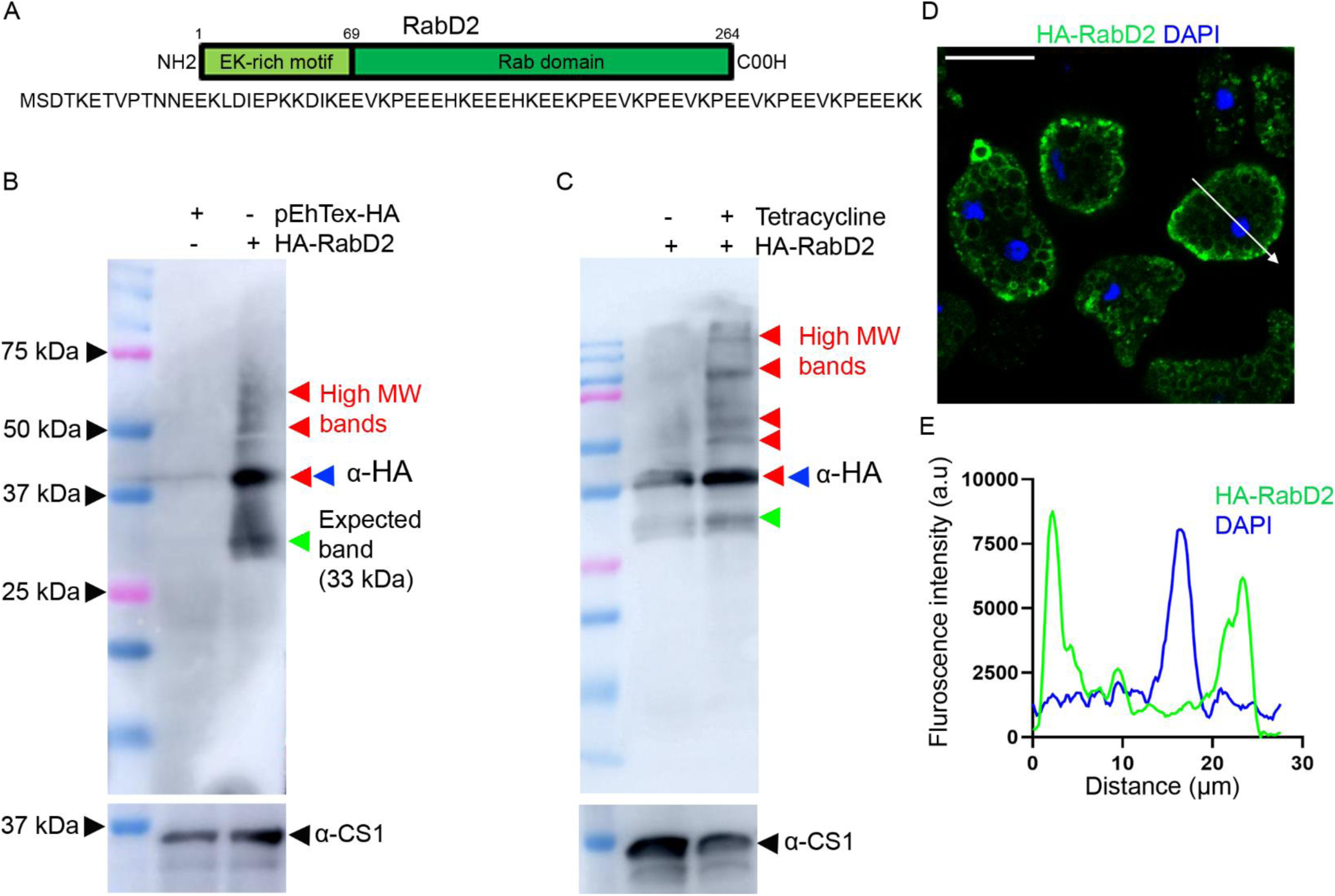
SDS-resistant amoebic RabD2 was observed at high molecular weight bands, and it is localized on variable size of vesicles and vacuoles. A. The domain organization of RabD2. The amino acid configuration-based domain mapping uncovers the N-terminal EK-rich motif and C-terminal Rab domain. B. The wild-type HA-RabD2 and pEhTex-HA (vector control) transfected trophozoites were incubated with tetracycline for 48 hours, and subsequently the trophozoite lysate was prepared and subjected to immunoblotting using anti-HA and anti-CS1 antibodies. The *red* arrowheads indicate the higher molecular weight bands, and the *blue* arrowhead indicates a highly intense, mid-modified, high molecular weight band compared to the hypothetical mass, which is depicted with a *green* arrowhead (the same color coding is followed in multiple blots). C. Wild-type HA-RabD2 expressing trophozoites in the absence (-) and presence (+) of tetracycline showed an increment in both expression and high molecular weight bands (in the presence of tetracycline). D. Wild-type HA-RabD2 expressing trophozoites were incubated on glass multi-well slides for 15 min at 37 °C and subsequently processed for immunofluorescence assay using anti-HA antibody and DAPI. Images were acquired using the confocal microscope. Scale bar, 20 µm. E. The graph shows fluorescence intensity of HA-RabD2 along the *white* arrow line depicted in Fig 1D, suggesting the distribution of RabD2 protein throughout the trophozoite.

### Endogenously, RabD2 self-associates, and it is sensitive to urea

We formerly observed the SDS-resistant high molecular weight species of RabD2 due to ectopic expression. To determine the nature of endogenous RabD2, amoebic RabD2 antibody was generated using the full-length RabD2 protein in rabbit and was validated (S2 Fig). Furthermore the validated antibody was used to confirm that the endogenous RabD2 is also SDS-resistant and reveals the high molecular weight bands (Fig 2A). The anomalous migration of proteins on SDS-PAGE could be a cause of post-translational modifications [29]. There are two common post-translational modifications, sumoylation (∼8 kDa) and ubiquitination (∼10 kDa), which lead to sequential increment of modified protein [30, 31]. Both the post-translational modification pathways are conserved in *E. histolytica* [32, 33]. The RabD2 sequence analysis also predicted seven potential sites (K24, 29, 43, 48, 53, 58, 63) with high confidence for sumoylation (ΨKXE/D (ψ: hydrophobic amino acid, K: target lysine, X: any amino acid, E and D: glutamic and aspartic acids, respectively) by SUMOplot (abcepta.com/sumoplot) and JASSA analysis tools [34]. Similarly, in the case of ubiquitination, ubiquitin is covalently linked with lysine (K) of the target protein through a three-step enzymatic cascade [35]. The 2-D08 and MG132 were widely used to selectively inhibit global sumoylation and ubiquitination, respectively [36, 37]. The untransfected amoebic trophozoites were incubated with 50 µM 2-D08 or 10 µM MG132 for 16 hours, and subsequently, it was identified that levels of RabD2 remained unchanged (Figs 2B and C). These results suggest that RabD2 neither undergoes sumoylation nor ubiquitination under physiological conditions. Few studies suggest that Rab GTPases undergo self-association or oligomerization in mammalian cells [38, 39]. However, the molecular basis of self-association in Rab GTPases and its functional relevance is not tested in any living system. To evaluate the self-association of RabD2 under steady-state conditions, wild-type amoebic trophozoites were grown at logarithmic phase, and the whole cell lysate was subjected to incubation with potent reducing agents like 10 mM dithiothreitol (DTT) and 10% β-mercaptoethanol (β-ME) for 30 minutes. We were not able to observe any remarkable difference in high molecular weight bands upon the treatment of reducing agents (S3 Fig). Next we hypothesized the possibility of ionic interactions that might be pivotal for the self-interaction of RabD2. Urea and guanidine hydrochloride (GnHCl) are chaotropic agents and commonly used as strong denaturants for proteins [40, 41]. The whole cell lysate was independently incubated with 6M urea and 9N GnHCl for 30 minutes. Our immunoblot result suggests that upon the treatment of urea, an intense band was observed at 30 kDa, which is likely to be the hypothetical size (monomeric version) of RabD2 (S3A Fig). Further, a similar experiment was performed at longer incubation time (16 hours) with urea and other reducing agents (S3B Fig). Surprisingly, we observed enrichment of RabD2 at expected hypothetical molecular mass (30 kDa) resulting from the decay of high molecular weight band (35 kDa). Finally, we wanted to study the effect of urea on the denaturation of endogenous RabD2 in a time-dependent manner. Trophozoites were lysed and incubated with urea for multiple time points. Our findings clearly suggest that the decay of high molecular weight bands, particularly 60-70 kDa, resulted in an intense 35 kDa band at 30 minutes.

**Fig 2.**
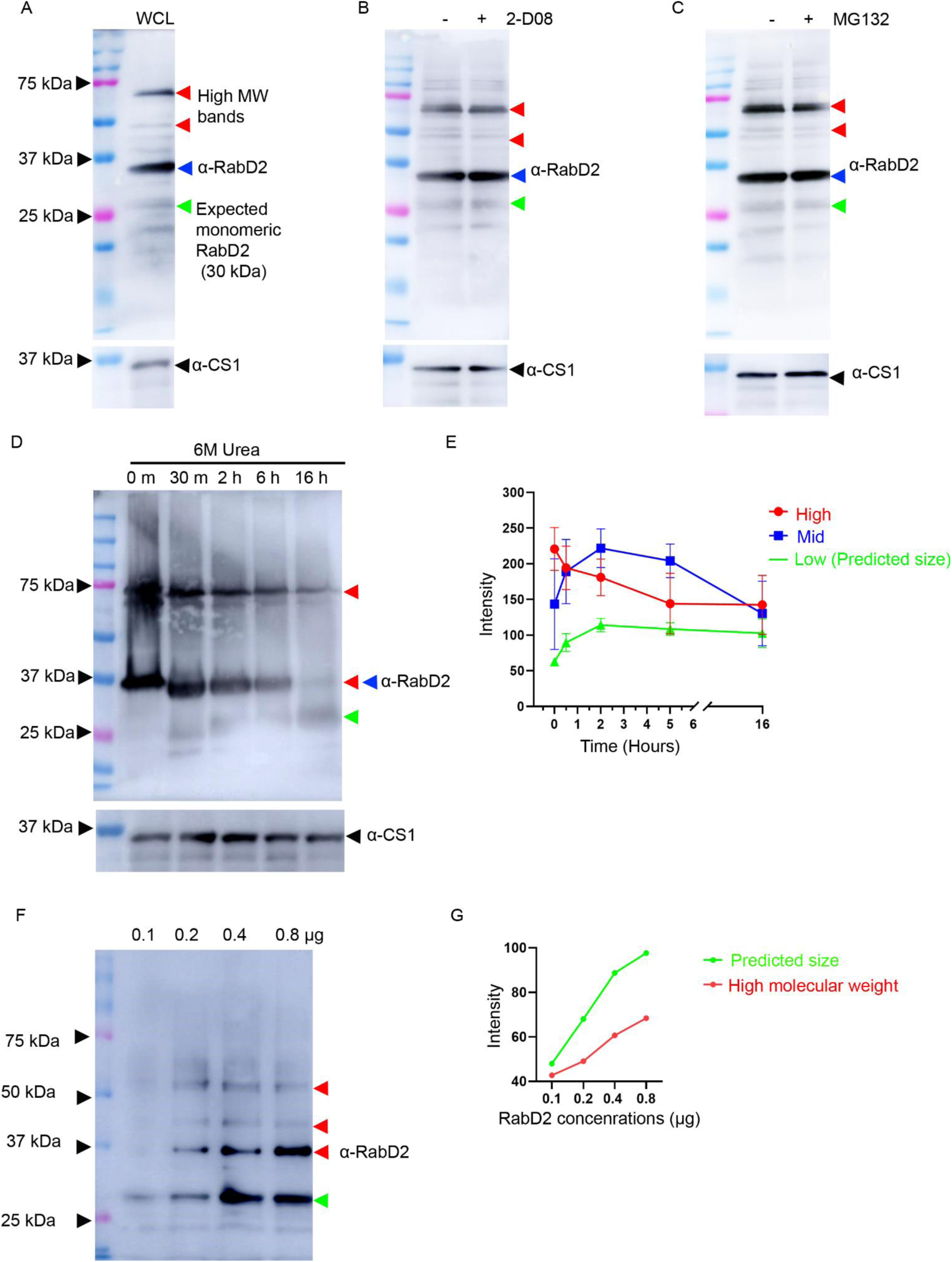
Endogenous RabD2 self-associates under physiological conditions. A. Wild-type trophozoites were lysed, and 50 µg of total protein was subjected to immunoblotting using anti-RabD2 and anti-CS1 antibodies. The expected band of RabD2 was faintly observed around 30 kDa (*green* arrowhead); additionally, intense high molecular weight bands (shown in *blue* and *red* arrowheads) were observed. B. C. Wild-type trophozoites were treated with 50 µM 2-D08 (B) or 10 µM MG132 (C) for 16 hours at 37 °C and then the lysate was subjected to immunoblotting using anti-RabD2 and anti-CS1 antibodies. D. Wild-type trophozoites lysate was incubated with 6M urea for different time points (0 minutes, 30 minutes, 2 hours, 6 hours, and 16 hours), separated on SDS-PAGE, and probed with anti-RabD2 and anti-CS1 antibodies. The *green* arrowhead indicates the hypothetical amoebic RabD2 band which is distinct from the mid-modified high molecular weight band (*blue* arrowhead) and the higher molecular weight bands (*red* arrowhead). E. The band intensity of the expected (30 kDa, *green*), highly intense mid-modified (35 kDa, *blue*), and the higher-order molecular weight bands (37-75 kDa, *red*) were quantified using the ImageJ software by maintaining uniformity in the area selected among all the lanes, thereby deducing the background. The graph represents the mean ± standard deviation (SD) calculated from three independent experiments. F. The purified amoebic RabD2 protein (0.1 - 0.8 µg) was separated by SDS-PAGE and electrophoretically transferred onto a PVDF membrane and probed with anti-RabD2 antibody. The *green* arrowhead shows a hypothetical molecular weight of the protein band around 30 kDa, and higher molecular weight bands are depicted in *red* arrowheads. G. Quantification of the band intensities for the expected (30 kDa) and the high-ordered bands (>30 kDa) was carried out using the ImageJ software. The graph represents the mean ± standard deviation (SD).

Eventually, the decay of the 35 kDa band into the monomeric version (30 kDa) of RabD2 in incremental time points was observed (Figs 2D and E). Further, the bacterially purified and proteolytically cleaved (PreScission Protease) RabD2 protein from the GST-RabD2 was employed in variable concentrations (0.1 - 0.8 µg) and subjected to immunoblotting. Our results showed that with increasing concentration of RabD2 protein, the high molecular weight bands appeared with enriched intensities at approximately 35 kDa, 45 kDa, and 60 kDa (Figs 2F and G). This result provides another line of evidence that RabD2 undergoes for high-ordered self-organization *invitro*. Collectively, our results showed that RabD2 self-associates potentially through ionic interactions and denatures in a urea-dependent manner.

### RabD2 self-associates through the atypical N-terminal glutamic acid-lysine (EK)-rich motif, which is controlled by the constitutively active and dominant negative mutants

We used a hypothetical protein sequence of RabD2 for *insilico* analysis and identified that RabD2 contains an EK-rich motif (1-69 amino acid residues) at the N-terminal (Fig 3A). To evaluate whether the N-terminal EK-rich motif is important for the self-association of RabD2, the EK-rich motif deletion construct (Δ69RabD2) was generated and ectopically expressed in amoebic trophozoites. The HA-Δ69RabD2 transfected trophozoites were selected, and their lysate was subjected to western blotting. Our results indicate that HA-Δ69RabD2 migrates only at the hypothetical molecular mass (approximately 24 kDa), and a significant reduction in the high-ordered self-association bands was observed (Figs 3B and C).

**Fig 3.**
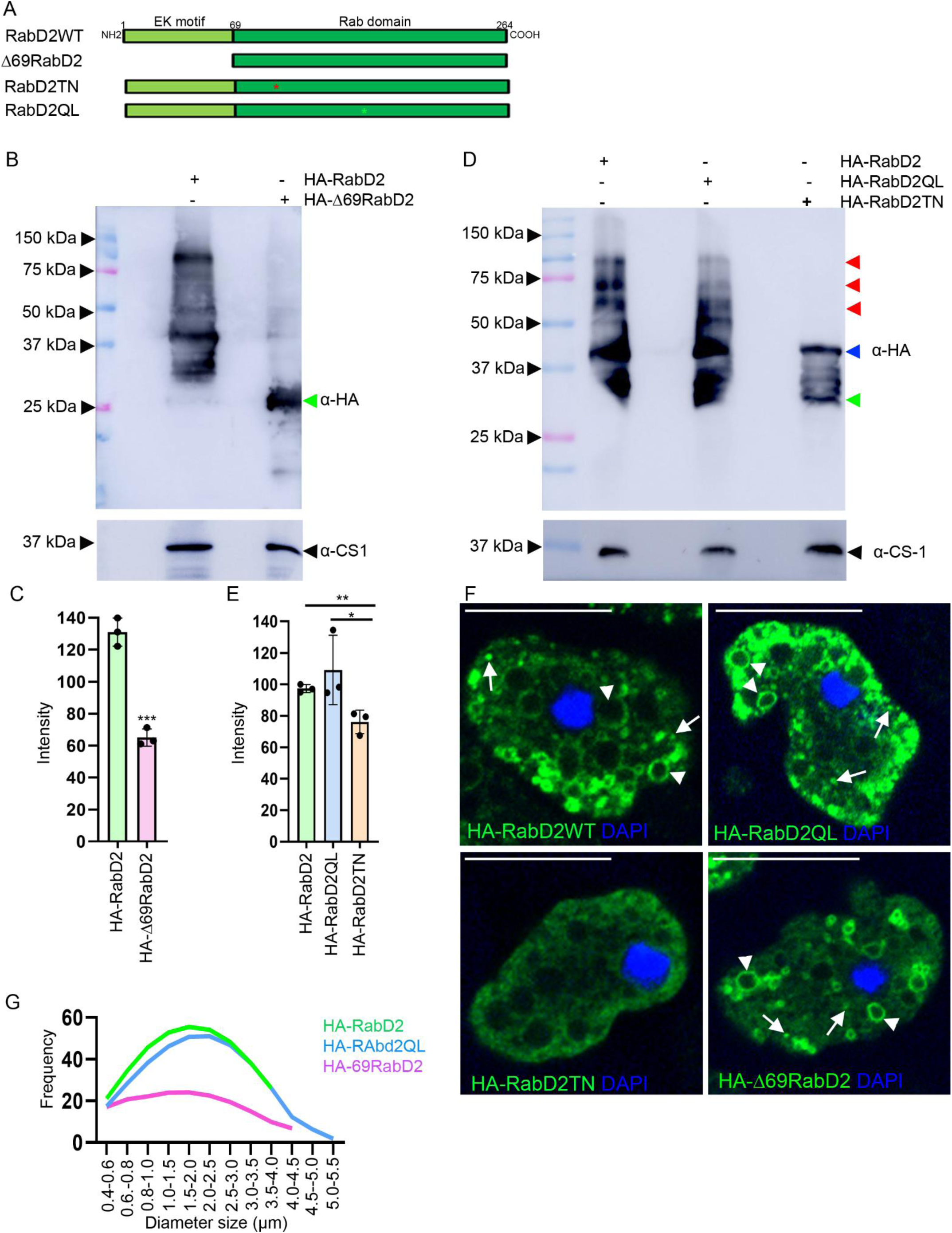
RabD2 self-association depends on the atypical N-terminal EK-rich motif, and it is controlled by the constitutively active and dominant negative mutants. A. Cartoon diagram of wild-type HA-RabD2 and its mutants used in the current study. The *red* asterisk indicates the site of the T91N mutation in HA-RabD2TN, and the *green* asterisk shows the site of the Q136L mutation in HA-RabD2QL mutants. B. The amoebic trophozoites expressing wild-type HA-RabD2 and truncated mutant HA-Δ69RabD2 were lysed, resolved by SDS-PAGE, and subjected to immunoblotting using anti-HA and anti-CS1 antibodies. Lane 2 shows a single band at approximately 24 kDa without any high molecular weight bands. C. The band intensities from >29 kDa were quantified using ImageJ. The bar graph represents the mean ± standard deviation acquired from the data analyzed from three independent experiments. Statistical significance from the represented graph is calculated with an unpaired student’s t-test (***P<0.001, p-value=0.0004). D. The cell lysates prepared from wild-type HA-RabD2, HA-RabD2QL, and HA-RabD2TN trophozoites were separated by SDS-PAGE and subjected to immunoblotting using anti-HA and anti-CS1 antibodies. A noteworthy reduction in the higher molecular weight bands was observed in the HA-RabD2TN mutant. E. Quantification of the band intensities >50 kDa was carried out using the ImageJ. The graph represents the mean ± standard deviation acquired from the data analyzed from three independent experiments. A drastic reduction in the high ordered bands was observed in HA-RabD2TN. Statistical significance from the represented graph is calculated with unpaired student’s t-test (**P<0.01, p-value=0.0096; *P<0.1, p-value=0.025). F. The amoebic trophozoites expressing HA-RabD2, HA-RabD2QL, HA-RabD2TN, and HA-Δ69RabD2 were incubated on glass slides for 15 min in BIS-33 medium at 37 °C. Trophozoites were fixed and processed for immunofluorescence assay with anti-HA antibody and DAPI. The *white* arrowheads (vacuoles) and *white* arrows (vesicles) show the distribution and localization of wild-type HA-RabD2 and its mutants. Scale bar, 20 µm. G. The histogram depicts the frequency and size distribution of RabD2 compartments in wild-type HA-RabD2 and its mutants. The dynamic size of vacuolar compartments was quantified using ImageJ, and the values obtained were normalized and plotted from three independent experiments. The graph represents the mean ± standard deviation and was fitted into a Gaussian distribution (300 vacuoles/30 trophozoites).

Next we investigated whether the nucleotide binding status of RabD2 controls the higher-order version of GTPase. We employed constitutively active mutant with defective GTP hydrolysis (HA-RabD2QL) and dominant negative mutant with defective GTP-binding (HA-RabD2TN). These standard mutations are established to show the nucleotide binding status of Rab GTPases in biochemical and cell biology studies [42, 43]. The RabD2 mutant (QL and TN) constructs were ectopically transfected, and its expression and subcellular localization was determined by immunoblotting and confocal microscopy. Strikingly, our immuno blot results showed that HA-RabD2TN has significantly reduced higher molecular weight bands compared to wild-type HA-RabD2 and HA-RabD2QL mutant (Figs 3D and E). These results suggest that RabD2 activity is remarkably contributing to the high-ordered organization of this protein. Our confocal imaging data indicates that the HA-RabD2 [diameter, 1.85±0.74 µm, n=364 compartments (including vesicles and vacuoles)/20 cells] and its mutants HA-RabD2QL (diameter, 2.07±0.9 µm, n=348 compartments/20 cells) and HA-Δ69RabD2 (diameter, 2.24±1.09 µm, n=182 compartments/20 cells) mostly localizes on variable sizes of vesicles and vacuoles (Fig 3F). Conversely, the self-association-deficient mutant HA-Δ69RabD2 localizes on variable sizes of vacuoles and shows a global reduction of 50% and 52.2% in total number of vacuoles compared to the wild-type RabD2 and RabD2QL mutant, respectively (Fig 3G). The localization of RabD2QL was similar to wild-type with a slightly high fraction of large-size (4.5-5.5 µm) vacuoles. Since the dominant negative mutation in Rab GTPases are known to interact with GDI (guanine nucleotide dissociation inhibitor), it is observed that their localization is mostly confined to the cytosol. Similarly, HA-RabD2TN is also localized in the cytosol, thereby attributing to its negligible appearance on the vacuoles and vesicles and thence excluded from this quantification. Altogether, our results suggest that the EK-rich motif and dominant negative mutant of RabD2 control the high-ordered self-association and subcellular distribution in amoebic trophozoites.

### RabD2 localizes on MVBs and controls its biogenesis

Our results indicate that the endogenous RabD2 localizes on large-size vacuoles suggestive to MVB-like structures, where discrete RabD2 vesicles (intraluminal vesicles, ILV) were observed within the large vacuoles (S4A Fig). Therefore, we sought to determine the localization of RabD2 with MVBs. Lysobisphosphatidic acid (LBPA) serves as a bona fide marker for ILVs within mature MVBs and is conserved from higher eukaryotes to *E. histolytica* [44, 45]. Hence, to test whether RabD2 is endogenously localized on MVBs, untransfected amoebic trophozoites were probed with anti-RabD2 and anti-LBPA. Interestingly, our confocal images clearly showed RabD2 vacuoles discreetly filled with LBPA (Fig 4A). The localization of RabD2 is observed to be abundant on the surface as well as within these vacuoles. Next, we performed transmission electron microscopy (TEM) to confirm that RabD2 localizes on MVBs. Indeed, our immunogold-labelled TEM micrograph showed that endogenously RabD2 localizes on the surface of MVB and ILV (Fig 4B). In eukaryotes, MVBs often originate from early endosomes, which are marked by the presence of Rab5 [46]. Therefore, HA-RabD2 expressing trophozoites were employed and probed with endogenous amoebic Rab5. Our confocal images clearly showed that RabD2 colocalizes with Rab5 on the surface of MVBs and ILVs (S4B Fig). Next, we sought to ask whether HA-RabD2 overexpression modulates the MVB abundance in the trophozoites. HA-RabD2 and pEhTex-HA trophozoites were subjected to TEM analysis. Our quantitative results suggest that HA-RabD2 expression showed a significantly high number of MVBs compared to pEhTex-HA (S5A and B Figs). These observations suggest that RabD2 could be potentially involved in MVB formation. Further, we asked whether RabD2 activity and its self-association are functionally important for MVB formation. HA-RabD2 and its mutants were probed with LPBA in steady-state conditions. We found that HA-RabD2, HA-RabD2QL, and HA-Δ69RabD2 mutants are localized on the surface of MVBs. Conversely, the dominant negative mutant HA-RabD2TN remained in the cytosol along with the diffused distribution of LBPA (Fig 4C). Next, we quantified the total number of LBPA-positive compartments and their varying size distribution in wild-type RabD2 and its mutants. Our quantification data suggest that HA-RabD2TN showed 17.69% and 52.5% reduction in total LBPA-positive compartments when compared with wild-type HA-RabD2 and HA-RabD2QL, respectively. Interestingly, we found that the size of LBPA compartments dramatically reduced in the HA-RabD2TN mutant (diameter, 1.86 ± 0.71 µm) when compared with wild-type HA-RabD2 (diameter, 2.73 ± 1.21 µm), HA-RabD2QL (diameter, 2.18 ± 1.25 µm), and HA-Δ69RabD2 (diameter, 2.22 ± 0.9 µm) mutants (Fig 4D). Altogether our results imply that RabD2 activity might be more important for LBPA association with MVBs. Therefore, to gain more details on the function of RabD2 in MVB biogenesis, RabD2 knockdown (KD) trophozoites were generated. The efficiency of RabD2 down regulation was measured in RabD2KD trophozoites compared with Trigger trophozoites (vector control). Our immunoblot quantification results showed significant reduction (around 50%) in overall RabD2 expression levels (Figs 4E and F). Further, RabD2KD trophozoites were probed for LBPA localization. Interestingly, our confocal micrographs clearly showed diffused LBPA, which is similar to that of HA-RabD2TN (Fig 4G). The diffused LBPA distribution suggests a defect in the classical MVB formation pathway. To gain a better understanding to MVB structures, the RabD2KD and Trigger trophozoites were subjected to TEM. Surprisingly, we observed that the RabD2KD trophozoites showed a large number of tiny vesicles and some of them are MVBs.). Interestingly, the large size dark MBVs with large ILVs were prominant in Trriger trophozoites compared with RabD2KD trophozoites (Fig 4H). In addtion, it was observed that in absence of RabD2, the size of ILVs reduced and showed close proximity with the MVB membrane. Upon quantification, we identified that RabD2KD trophozoites showed a significant reduction in the diameter of MVBs (Fig 4I), which implies an improper maturation of MVBs. Altogether, our results suggest that RabD2 localizes on MVBs and controls their biogenesis.

**Fig 4.**
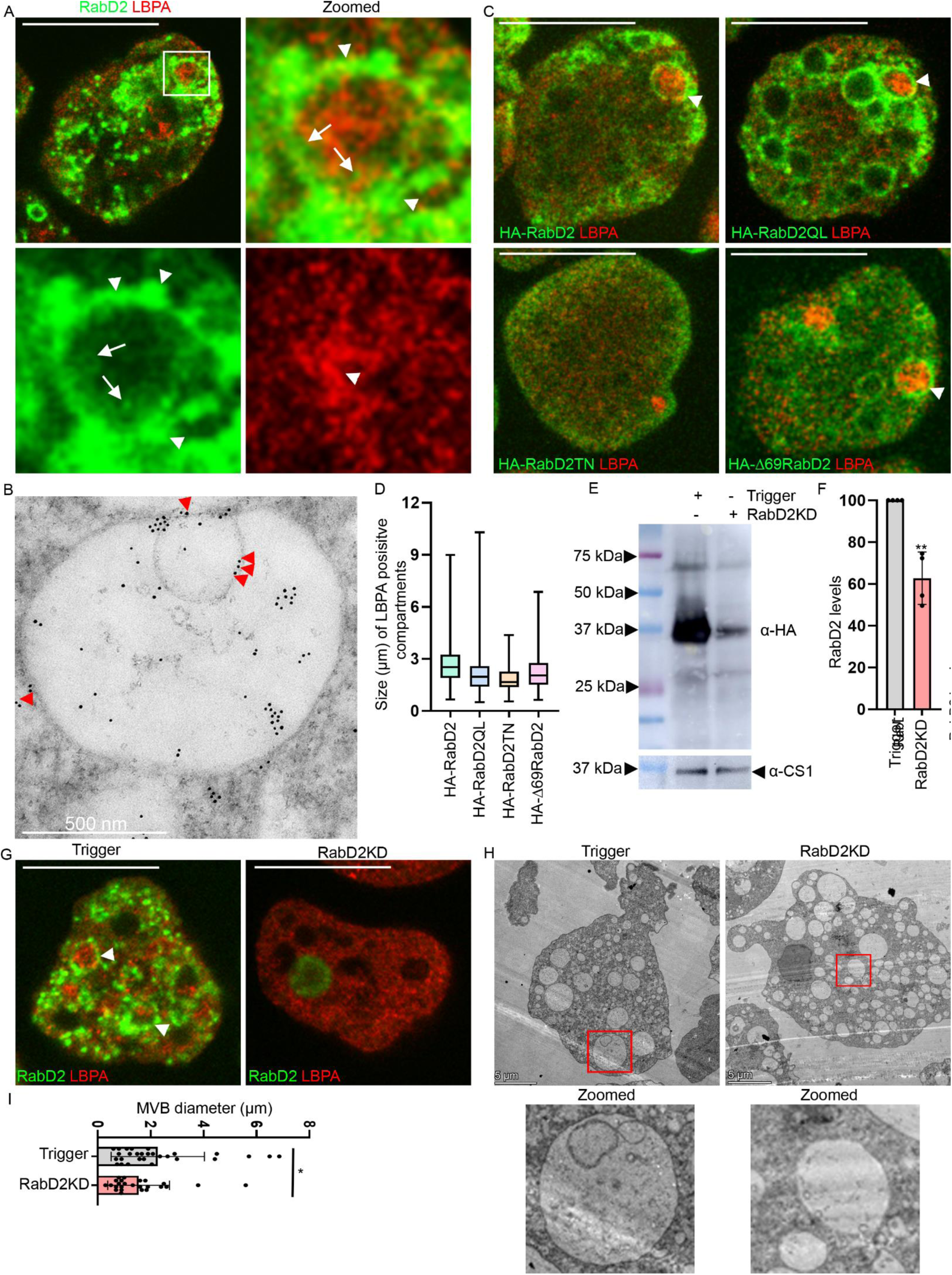
RabD2 localizes on MVBs and controls its biogenesis. A. Endogenous subcellular localization of RabD2 in *E. histolytica*. Wild-type trophozoites were incubated on glass slides for 15 min in BIS-33 medium at 37 °C. Trophozoites were fixed and processed for immunofluorescence assay with anti-RabD2 and anti-LBPA antibodies. The square box represents the zoomed panel showing LBPA-filled vacuole decorated with RabD2 vesicles. The *white* arrowheads (in the *green* panel) indicate the localization on the surface of MVB, and the *white* arrows (in the *green* panel) show the localization within the MVBs. The *white* arrowhead (*red* panel) indicates LBPA puncta. Scale bar, 20 µm. B. Transmission electron micrograph featuring immunogold labelling with anti-RabD2 antibody and probed with 10 nm gold-conjugated secondary antibody in wild-type trophozoites. The *red* arrowheads show the localization of RabD2 on MVB and ILV. Scale bar, 5 µm. C. The trophozoites expressing wild-type HA-RabD2, HA-RabD2QL, HA-RabD2TN, and HA-Δ69RabD2 were incubated on glass slides for 15 min, fixed and processed for immunofluorescence assay with anti-HA and anti-LBPA. The *white* arrowheads show the distribution, localization and association of wild-type HA-RabD2 and its mutants with LBPA. Scale bar, 20 µm. D. The total number and size of the LBPA compartments were quantified using ImageJ. The graph represents the mean ± standard deviation of LBPA compartments acquired from the pooled data of three experiments (n = 300 cells). E. The RabD2KD and Trigger trophozoites were lysed and subjected to western blotting using anti-RabD2 and anti-CS1 antibodies. F. The band intensities were quantified using ImageJ. The graph represents the mean ± standard deviation acquired from the data analyzed from three independent experiments. The statistical significance was calculated using an unpaired Student’s t-test (**P<0.01, p-value=0.001). G. The Trigger and RabD2KD trophozoites were subjected to immunofluorescence and were probed with anti-RabD2 and anti-LBPA antibodies. The *white* arrowheads indicate the association of RabD2 vacuoles with LBPA. Scale bar, 20 µm. H. The Trigger and RabD2KD trophozoites were subjected to TEM; the *red* square box represents the zoomed panels highlighting the individual MVBs and their ILVs, emphasizing the morphological differences. Scale bar, 5 µm. I. The number and diameter of MVBs were quantified using ImageJ. The graph represents the mean ± standard deviation. The statistical significance was calculated using unpaired Student’s t-test (*P<0.1, p-value=0.047) (n=25 MVBs/10 cells).

### RabD2 upregulation causes global ubiquitination, degradation of the heavy chain of GalNAc lectin (Hgl), and inhibition of adherence

It is very well established that specific sets of cargoes and receptors undergo MVB-dependent protein degradation in eukaryotes [47–49]. Therefore, we looked for the ubiquitination status in amoebic trophozoites upon the ectopic expression of RabD2. Surprisingly, we have observed that the overall ubiquitination is enhanced in RabD2 expressing trophozoites compared to uninduced RabD2 and pEhTex-HA trophozoites (S5C Fig). Further, we showed that the constitutively active HA-RabD2QL mutant exhibits high levels of ubiquitination compared to the dominant negative HA-RabD2TN and HA-Δ69RabD2 mutants (Figs 5A and B). These results indicate that RabD2 expression is sufficient to induce ubiquitination, which potentially controls the degradation of specific receptors in amoebic trophozoites. The GalNAc lectin, a tetrameric complex constituting the Hgl, is required for host cell adhesion, cytotoxicity, and complement resistance [50–52]. Therefore, we asked how RabD2 expression and its activity impact the levels of Hgl in amoebic parasites. Interestingly, we found that the HA-RabD2QL mutant significantly causes down regulation of Hgl, which is 52.8% and 54.7% when compared to the HA-RabD2TN and HA-Δ69RabD2 mutants, respectively (Figs 5C and D). These results suggest that both activity and the EK-rich motif of RabD2 are involved in the down regulation of Hgl. Therefore, we asked whether the down regulation of Hgl controls the adherence of trophozoites to the host cells. The wild-type HA-RabD2 and its mutant trophozoites were incubated with Chinese hamster ovary (CHO) cells for adhesion (S6 Fig). Interestingly, our quantification results showed that HA-RabD2QL expression significantly reduced its adherence to the CHO cells, which is 71.22% and 61.01% when compared with HA-RabD2TN and HA-Δ69RabD2, respectively (Fig 5E). These results raise the possibility that HA-RabD2QL mutants poor adherence to CHO cells is due to the relatively low level of Hgl at the cell surface. The plasma membrane levels of Hgl were observed in previous studies using the non-permeabilized trophozoites [53, 54]. Further, cell surface-associated Hgl was investigated in wild-type HA-RabD2 and its mutants. Our imaging results showed that plasma membrane abundance of Hgl dramatically reduced in HA-RabD2QL and HA-RabD2WT compared with HA-RabD2TN and HA-Δ69RabD2 mutants (Figs 5F and G). Similarly, our quantification data indicates that the percentage of trophozoites with plasma membrane localization of Hgl was significantly reduced in HA-RabD2 WT and HARabD2QL mutants (Fig 5H). Altogether, our results clearly showed that the activity of RabD2 causes global ubiquitination that ultimately downregulates the plasma membrane levels of Hgl and reduces the adherence of trophozoites to the host cells.

**Fig 5.**
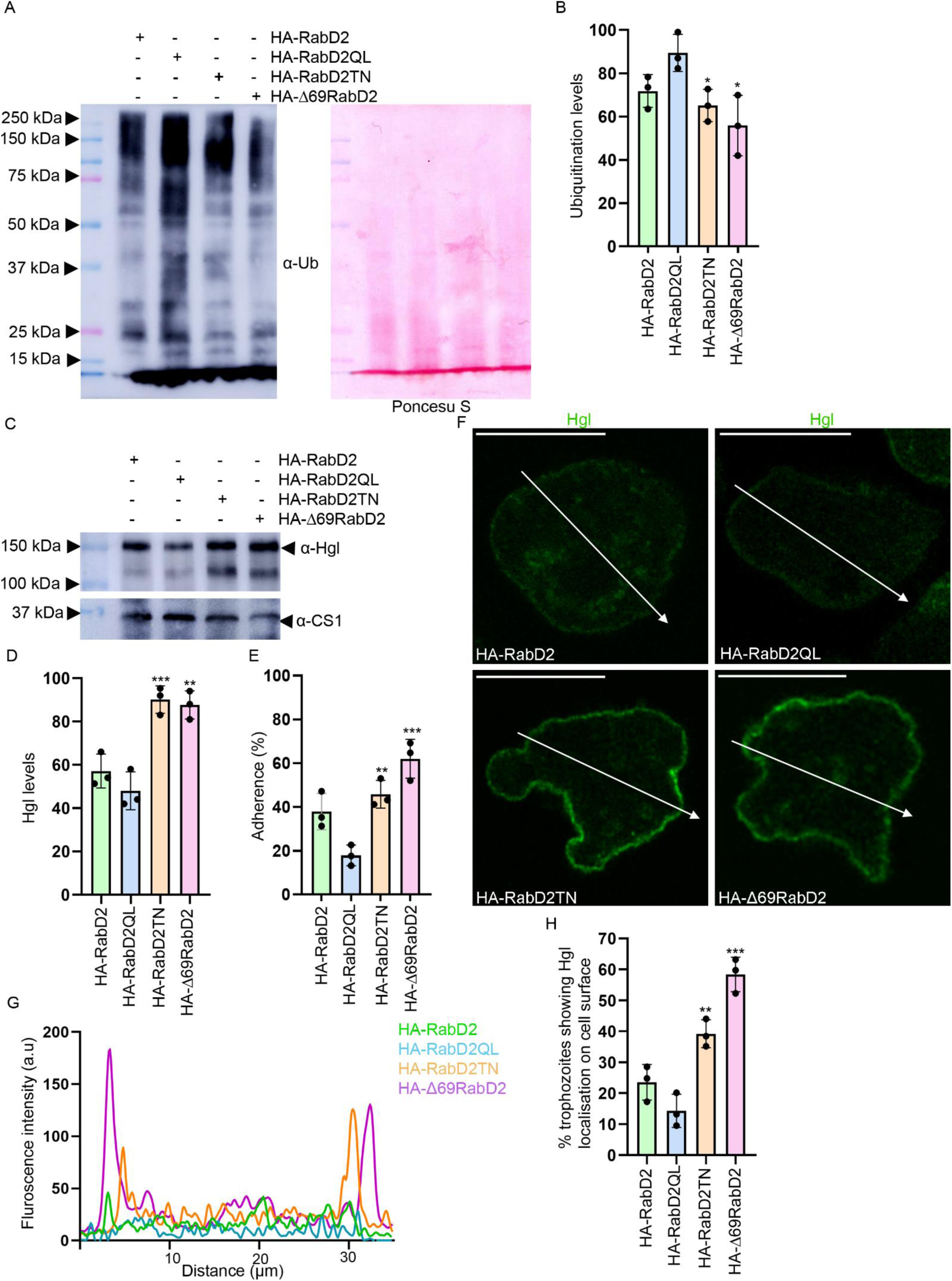
RabD2 expression upregulates the global ubiquitination leading to down regulation of Hgl and adherence to host cells. A. To evaluate the ubiquitination status in wild-type HA-RabD2 and its mutants. HA-RabD2WT and its mutants (HA-Rab2QL, HA-RabD2TN, and HA-Δ69RabD2) were lysed and subjected to immunoblotting with anti-ubiquitin and anti-CS1 antibodies. The left panel shows the Ponceau S staining for the similar blot as an evaluation of loading control. B. The relative signal intensities from the anti-ubiquitin blot were quantified and plotted using ImageJ. The bar graph represents the mean ± standard deviation (SD) calculated from three independent experiments. The statistical significance was calculated between RabD2QL, RabD2TN and Δ69RabD2 using an unpaired Student’s t-test (*P<0.1, p-value=0.021). C. Wild-type HA-RabD2 and its mutants (RabD2QL, RabD2TN, and Δ69RabD2) trophozoites were lysed and subjected to immunoblotting with anti-Hgl and anti-CS-1 antibodies. D. Quantification of anti-Hgl blot intensities was done using imageJ. The bar graph represents the mean ± standard deviation (SD) calculated from three independent experiments. The statistical significance was calculated between RabD2QL and Δ69RabD2 using an unpaired Student’s t-test (**P<0.01, p-value=0.0025, **P<0.01, p-value=0.0032). E. Wild-type HA-RabD2 and its mutants (HA-Rab2QL, HA-RabD2TN, and HA-Δ69RabD2) were co-incubated with CHO cells (1:20 ratio) for 90 minutes and then imaged. The quantitative adherence of amoebic trophozoites to CHO cells was manually quantified. Adherence was considered positive when more than two CHO cells were adhered to the amoebic trophozoite. Data represents mean ± SD from three independent experiments. The statistical significance was calculated between RabD2QL, RabD2TN, and Δ69RabD2 using an unpaired Student’s t-test (***P<0.001, p-value=0.0006; **P<0.01, p-value=0.0036) (n=75 cells/construct). F. Wild-type HA-RabD2 and its mutants (HA-RabD2QL, HA-RabD2TN, and HA-Δ69RabD2) trophozoites were incubated under steady state for 30 min at 37 °C. The trophozoites were fixed with 4% paraformaldehyde, blocked with 5% FBS, and probed with anti-Hgl antibody. Scale bar, 20 µm. G. The graph shows the fluorescence intensities of wild-type HA-RabD2 and its mutants (HA-RabD2QL, HA-RabD2TN, and HA-Δ69RabD2) trophozoites along the *white* arrow line depicted on Fig 5F showing the membrane localization of Hgl. The graph represents the mean ± standard deviation. H. Quantification of trophozoites showing localization of the plasma membrane Hgl. The number of trophozoites with plasma membrane Hgl was calculated from the three independent experiments. The graph plotted represents the mean ± S.D. Statistical significance was calculated between RabD2QL, RabD2TN, and Δ69RabD2 using an unpaired two-tailed Student’s t-test. (***P<0.1, p-value=0.0006) (n=120 cells/experiment).

## Discussion

This study presents three major findings. First, we identified the presence of a novel EK-rich motif at the N-terminal of the conserved Rab domain that participates in self-association.

Second, RabD2 localizes on MVBs and controls their biogenesis. Third, we found that RabD2 self-association and GTPase activity regulate global ubiquitination accompanied with the down regulation of the plasma membrane receptor Hgl.

Rab GTPases are ubiquitous in eukaryotes and are crucial for the transport of cargoes between compartments via vesicular budding, fusion, and docking [13]. The amoebic RabD2 protein is relatively longer than the conventional small Rab GTPases because of its EK-rich motif at the N-terminal of Rab domain. The presence of this motif is unique in the Rab proteins. In mammals, a subset of Rab GTPases is of higher molecular weight compared to classical small Rab GTPases. These high molecular weight Rab GTPases, known as large Rab GTPases, consist of an N-terminal EF-hand domain (EFD), a coiled-coil domain (CCD), and a C-terminal conserved Rab G domain. Rab GTPases 44, 45, and 46 belong to this class in mammals [25]. It has been demonstrated that the N-terminal domains are required for the self-association of these GTPases under physiological conditions [38, 39]. Conversely, the Rab40 family of proteins also contains an additional region at C-terminal, the suppressor of cytokine signalling (SOCS) box [55]. Our study identified that the novel N-terminal EK-rich motif (potential disordered region) is crucial for controlling the self-association of RabD2. Surprisingly, we observed that RabD2 forms high-order oligomers that are resistant to SDS and other denaturants. Additionally, we found that the chaotropic agent urea denatures the protein into its monomeric version of RabD2. Both GnHCl and urea are chaotropic agents that disrupt the non-covalent bonds within the protein’s tertiary structure [56]. Previously, it has been shown that urea and GnHCl are structurally similar and denature proteins by forming hydrogen bonds with the NH_2_-group and COOH-group of peptides [57]. In contrast, lysozyme stability sharply decreases upon GnHCl treatment, whereas in the presence of urea, the protein remains folded and stable [58]. Urea solubilizes and denatures proteins by disrupting the non-covalent interactions that stabilize their secondary and tertiary structures. Urea denatures proteins through both direct and indirect mechanisms [59, 60]. The direct mechanism involves hydrogen bonding of urea with polar amino acids of the protein, leading to weak intermolecular bonds and interactions between amino acid residues. Urea can also compete with the protein for water molecules, reducing solvation and hydration of the protein and exposing its hydrophobic core. The indirect mechanism involves altering the solvent environment by urea, affecting the thermodynamics and kinetics of protein folding and unfolding. Urea can change the structure and hydrodynamics of water, making it more like a non-polar solvent, which favours the denatured state of the protein. Collectively, it can be speculated that thermodynamic parameters, overall amino acid composition, and peptide charge in EK-rich motifs confer to maintaining protein stability in the presence of different chaotropic agents. The N-terminal region of RabD2 provides a negatively charged side chain of glutamic acid (E) and a positively charged side chain of lysine (K) that may contribute to salt-bridge formation. It has been shown that the ionic interactions between the positive (aspartic acid, glutamic acid) and negative (lysine, histidine) side chains of charged amino acids are crucial for structural integrity and function [61, 62]. Interestingly, we observed that high-ordered self-association of RabD2 is controlled by an EK-rich motif and dominant-negative mutant (GDP-bound version). This result raises the possibility that G-domain and EK-rich motif interaction may contribute to the GTPase activity of RabD2. It has been shown that ARF (ADP-ribosylation factor) protein 1, GTP binding allows conformational changes at the N-terminal amphipathic helix that are required for membrane association [63]. Future study might require checking whether EK-rich motif of RabD2 have specific preferences for lipids for membrane recruitment and function similar to ARF proteins [64].

MVBs are ancient organelles and are found across various eukaryotic lineages [6]. The direct role of Rab GTPases in MVB formation is poorly studied from lower to complex eukaryotes. The mammalian Rab11, Rab27a, and Rab27b have been shown to be involved in MVB fusion and exosome biogenesis, although their functional roles are cell type-specific [17, 19]. We identified that RabD2 localizes on LBPA-positive compartments and MVBs through confocal and TEM imaging. Previously, RabD2 was identified as an interacting partner of endogenous EhVps23 in immunoprecipitation based proteome studies. EhVps23, an orthologue of yeast Vps23 and mammalian TSG101, undergoes ubiquitination and is recruited to MVBs and phagosomes [10]. Our results corroborate previous studies where EhVps23 localizes on MVBs. Recent studies showed that the EhVps35 C, a member of a retromer complex tie-up with ESCRT machinery proteins [65]. RabD2 was also identified in EhVps35 C interactome along with ESCRT-I and II members. We also observed that overexpression of RabD2 causes significant upregulation of MVBs and are localized at the abscission site of ILV where EH domain-containing protein (EHD1) is localized [27]. Further, it would be interesting to study how RabD2 crosstalks with ESCRT machinery proteins and how membrane lipids specificity controls the organization of MVBs. Interestingly, while ESCRT complexes I, II, III, and III-associated are ubiquitous and conserved in eukaryotic lineages. ESCRT 0 members appear to be missing in this lineage and have specific requirements to initiate the process. It has been shown that EhTom1, similar to *D. discoideum* DdTom1 (VHS and GAT), interacts with TSG101 and facilitates the sorting of ubiquitinated proteins onto MVBs [65]. Future study is required to establish the direct link between ESCRT and retromer complex proteins with RabD2.

The crosstalk between MVBs and ubiquitination is crucial for regulating protein sorting and degradation within the cells [49]. We have shown that RabD2 ectopic expression causes upregulation of global ubiquitination of proteins in amoebic trophozoites. This raises the question of how RabD2 overexpression causes this. First, it can be predicted that RabD2 activates ubiquitin conjugation machinery proteins. It has been found that the SOCS box (non-Rab region of Rab40c), an evolutionarily conserved LPLP motif, is required for the binding to Cullin5 that contributes to the formation of ubiquitin E3 ligase complex, which regulates protein ubiquitination [55, 66]. Second, RabD2 may recruit ESCRT complex proteins to effectively target ubiquitinated proteins on endosomes. The RabD subfamily proteins are conserved in protozoa and plant species and may have been evolutionarily lost in mammals. In *Arabidopsis thaliana*, RabD1 and RabD2 GTPases localize on Golgi/TGN and control the trafficking to specific endosomes [67]. We can also predict that RabD2 vesicles may originate from the Golgi and traffic the cargoes to endosomes. Interestingly, it has been shown that plant ESCRT machinery proteins transport cargoes to the endosome from the TGN. Third, RabD2 activity in the late endocytic pathway may trigger receptor proteins to be directed towards lysosome fusion and degradation. In mammals, Rab7 is required for the biogenesis of MVBs and degrades the cargoes through lysosomal fusion [16]. Rab7b, on the other hand, has been shown to promote the lysosomal degradation of Toll-like receptor 4 (TLR4) [68]. Interestingly, we cannot rule out the possibility that RabD2 is also involved in the proteasomal degradation pathway, as it has been shown that ubiquitinated proteins are also degraded through this path. The contrasting observation is suggesting that proteasomal function and ubiquitination levels cannot be correlated in amoebic parasites. Because proteasome function is highly upregulated in the trophozoites, encystation and ubiquitination levels remain fluctuating in the same process [69]. Further, it would be interesting to identify the RabD2 downstream effector molecules and their roles in the above possibilities.

The upregulation of amoebic RabD2 was specifically identified upon interaction with human colon explants and under oxidative stress conditions [20]. These observations suggest that host tissue interaction potentially causes a stress response leading to global reprogramming in the trophozoites. This situation may allow the trophozoites in the host environment to adapt by down regulating their own proteins through ubiquitination. Recently, antibodies to *E. histolytica* ubiquitin have been identified in patients with invasive amoebiasis [70]. In addition, it has been shown that the Eh ubiquitin also reduces robust immune response in animal models [71]. It is possible that the trophozoites ubiquitinated proteins containing MVBs fuse with the plasma membrane and exocytose extracellular vesicles (EVs) into the host environment. Interestingly, it has been observed that *E. histolytica* EVs downregulate the human neutrophil respiratory burst and NETosis [72]. It can be speculated that EV secretion from the trophozoites is potentially involved in immune evasion. In the current study, down regulation of Hgl was identified in the constitutively active mutant RabD2QL. Conversely, accumulation of Hgl was observed in the catalytically inactive and self-association deficient mutants. We believe that with increased MVB formation, more Hgl is sequestered into ILVs within MVBs. These MVBs have two fates: either fusing with lysosomes for degradation or fusing with the plasma membrane to release exosomes. Previously, it has been shown that Hgl is constitutively degraded through the endo-lysosomal pathway [54]. Interestingly, a recent study found a 5.8-fold increase in Hgl abundance in the EV population upon the interaction with host neutrophils [72]. It could be possible that the down regulation of membrane receptors is considered a double-edged sword in which trophozoites reduce adherence to tissue, favouring evasion from the host immune system. This might be a potential mechanism for the parasites long-term survival, replication, and establishing the infection in the host. We cannot rule out the other possibility for constitutive exocytosis of Hgl that could lead to global depletion of Hgl in the trophozoites and modulate immune surveillance. Interestingly, we found that plasma membrane abundance of Hgl and adherence of trophozoite is reduced in wild-type RabD2 and RabD2QL mutant. In this context the recent study has highlighted the tetraspanins interaction with Hgl and its involvement in cargo selection and its packaging onto MVBs in *E. histolytica* [73]. However, tetraspanins 12 and 13 overexpression significantly reduced the adherence and upregulated the proteases secretion. It is noteworthy to check the involvement of RabD2 in the secretion of extracellular vesicles from *E. histolytica* against the host cell killing and immune modulation.

Collectively, we have identified that RabD2 self-associates and controls the down regulation of Hgl, leading to decreased adherence of the trophozoites to the host through the MVB pathway (Fig 6).

**Fig 6.**
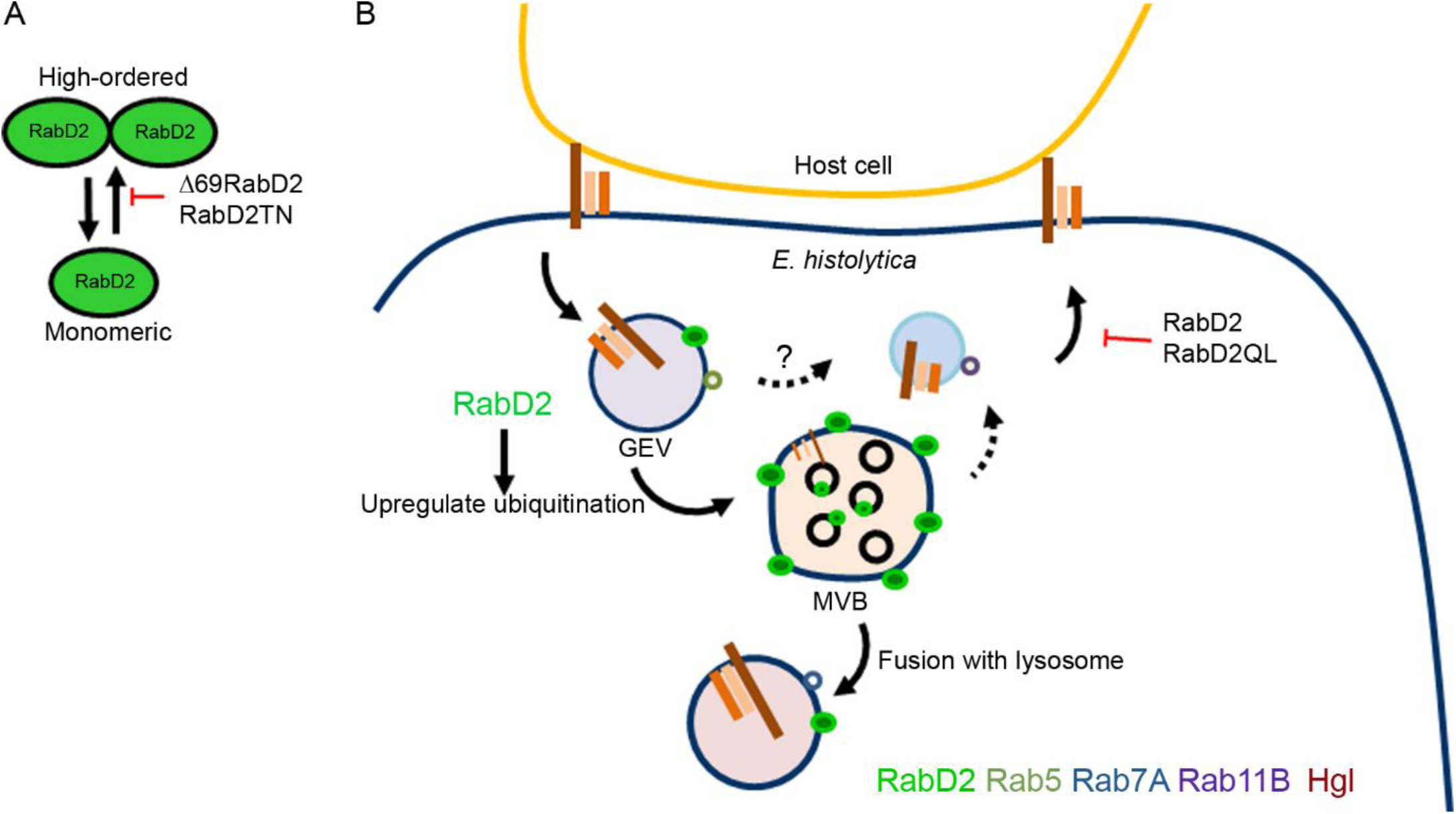
The proposed model for amoebic RabD2-mediated MVB formation regulates the trophozoite adherence to the host cell. A. RabD2 self-association is controlled by the N-terminal EK-rich motif and dominant negative mutant. B. The overexpression of RabD2 causes upregulation of ubiquitination, ultimately leading to the down regulation of Hgl. The data from previous studies suggest that Hgl is internalized from the plasma membrane and transported to lysosome-like compartments through giant endocytic vacuoles (GEVs positive for Rab5 and Rab7A) or recycled (via Rab11B) to the cell surface [54]. RabD2 recruits at the Rab5-positive endosomes and facilitates the formation of MVBs. Subsequently, Hgl is either trafficked to an acidic lysosomal compartment or exocytosed via MVBs. The self-association of RabD2 may be required for Hgl trafficking to the plasma membrane. In the absence of RabD2 self-association, Hgl may get stuck in MVBs and fail to degrade.

## Materials and Methods

### Cell culture

*Entamoeba histolytica* HM-1: IMSS trophozoites were cultured under axenic conditions in BIS-33 medium supplemented with 15% (v/v) adult bovine serum (RM10913, Himedia Labs, India), 2% (v/v) Diamonds vitamins, 100 units penicillin, 100 µg/mL streptomycin, 0.25 µg/mL and amphotericin B (15240062, Gibco, USA) and the amoebic trophozoites were maintained at 35.5°C [74].

### cDNA pool preparation and cloning

Logarithmic phase trophozoites were used to extract the total amoebic RNA using PureLink™ RNA Mini Kit (12183020, Invitrogen, USA) as per manufacturer’s instructions. The obtained total RNA was used in the synthesis of cDNA with a High-Capacity RNA-to-cDNA™ Kit (4387406, Invitrogen, USA). 50 ng of the pooled cDNA together with amoebic RabD2 (EHI_164900) primers (forward primer 5’gcacccgggATGAGTGATACTAAAGAGACTGTTCC3’ and reverse primer 5’gacctcgagTTAACAACATCCTCCTTTCTTTT3’) were used in PCR amplification using Phusion® High-Fidelity PCR master mix (M0531S, New England Biolabs, USA). The amplified products were electrophoresed on 1% (w/v) agarose gel and the predicted molecular weight band (approximately 792 bp) was excised and digested with *SmaI* and *XhoI* restriction enzymes (R0141S, R0146S New England Biolabs, USA). The pEhTex-HA plasmid was also digested with aforementioned enzymes and was subjected to ligation with the RabD2 PCR product using T4 DNA ligase (M0202S, New England Biolabs, USA). The ligated product was transformed to *E. coli* DH5-α, and recombinant clones (pEhTex-HA-RabD2) were identified and confirmed using unidirectional Sanger sequencing.

### Site-directed mutagenesis and generation of deletion mutants

The RabD2 mutant primers were designed using the NEBaseChanger tool (nebasechanger.neb.com). To create the mutant constructs, 25 ng of HA-RabD2 plasmid was amplified using the constitutively active RabD2Q136L (RabD2QL) primers (forward primer 5’TACAGCTGGACTGGAACGTTTTG3’ reverse primer 5’TCCCAAATTTGAAGTTTAATATG3’) and dominant negative RabD2T91N (RabD2TN) primers (forward primer 5’TGTTGGTAAAAACTGTTGTATGAATAGATATGTTAG3’ reverse primer 5’GAAGACTCTCCTACCATAATAAC3’) using Q5 Site-directed mutagenesis kit (E0554S New England Biolabs, USA) as per manufacturer’s instructions. The deletion construct Δ69RabD2 was generated by PCR amplification using these primers (forward primer 5’gcacccgggAATGAATCTAATGAACAAC3’ reverse primer 5’gacctcgagTTAACAACATCCTCCTTTCTTTT3’).

### Electroporation and selection of stable expressing trophozoites

*E. histolytica* trophozoites at mid-logarithmic growth phase were harvested by replacing the culture media with cold 1X phosphate saline buffer (PBS) pH 7.4 and placing the tube on ice for 10 minutes (to detach cells). The cell suspension was then centrifuged at 1000 rpm for 5 minutes and the pellet was suspended in ice-chilled incomplete cytomix buffer (10 mM K₂HPO₄/KH₂PO₄ (pH 7.6), 25 mM HEPES (pH 7.4), 120 mM KCl, 0.15 mM CaCl₂, 2 mM EGTA, and 5 mM MgCl₂) subsequently centrifuged at 900 rpm for 5 minutes. The pellet was resuspended in complete cytomix buffer (incomplete cytomix buffer supplemented with 10 mM reduced glutathione and 4 mM ATP) along with 80 µg of plasmid DNA in a 0.4cm electroporation cuvette (1652088 Biorad, USA) with modifications [75]. The trophozoites were then electroporated at 500V voltage, 500µF capacitance, and time constant infinity (∞) using Gene Pulser Xcell Eukaryotic System (Biorad, USA). Immediately the contents in the electroporation cuvette were transferred into warm complete culture media (BIS-33 supplemented with 15% adult bovine serum and 2% Diamonds vitamins) without antibiotics. The stable transfectants were screened using G418 (A1720, Sigma-Aldrich, USA) selection, which involves gradual increment of G418 from 2 µg/ml to 6 µg/ml with regular change of media for 48 hours over a period of 12-15 days and the stable trophozoites were maintained at 6 µg/ml G418. Before every experiment performed, the stable trophozoites were shifted to complete BIS-33 medium supplemented with 20 µg/ml G418 and 5 μg/ml tetracycline to induce the expression of the target gene for at least 36-48 hours.

### Indirect immunofluorescence assay and confocal microscopy

Trophozoites at mid-logarithmic phase were washed with cold 1X PBS and harvested by centrifugation at 1000 rpm for 5 minutes. The pellet was then suspended in warm BIS-33 medium and transferred onto a clean multitest 8-well slide (6040805, MP Biomedical, USA), and incubated at 37°C in a water bath for 20 minutes. The adhered trophozoites were fixed with 4% (w/v) paraformaldehyde (158127, Sigma-Aldrich, USA) for 15 minutes, followed by a wash with 1X PBS and then permeabilized with 0.1% (v/v) Triton X-100 (in 1X PBS) for 9 minutes. Blocking is carried out in 5% (v/v) fetal bovine serum (26140079, Thermo Fisher Scientific, USA) prepared in 1X PBS (blocking solution) for 30 minutes. Trophozoites were incubated with specific primary antibodies; anti-HA monoclonal antibody (sc-7392, Santa Cruz Biotechnology, USA) (1:150) or anti-RabD2 (1:500) or anti-LPBA (Z-PLBPA, Echelon Biosciences) (1:100) or anti-Hgl (1:100, 3F4 and 7F4) or anti-Rab5 (1:50) for 90 minutes subsequently three washes with blocking solution were given. The secondary antibodies conjugated with Alexa Fluor 488, 568, and 647 (Alexa Fluor 488 anti-mouse, A11029; Alexa Fluor 488 anti-rabbit, A11034; Alexa Fluor 568 anti-mouse, A11031; and Alexa Fluor 647 anti-mouse, A21235; Invitrogen, USA) were prepared in blocking solution (1:500) and incubated for 60 minutes in the dark. To probe the DNA, DAPI (10236276001, Sigma-Aldrich, USA) (1:1000) is co-incubated with secondary antibodies. Finally, mounting of trophozoites with coverslips was done using 5 µl of Prolong Diamond Antifade (P36970, Invitrogen, USA), and the slides were dried at room temperature overnight and employed for microscopy.

The image acquisition was done using the LSM 900 laser scanning confocal microscope (Carl Zeiss, Germany) and the SP8 laser scanning confocal microscope (Leica Microsystem, Germany) equipped with multiple laser lines (405, 488, 561, and 640 nm). The 63X oil immersion objective (with 1.4 NA) was used, and all the images were acquired at 8-16 bit resolution, 512x512 or 1024x1024 frames, with a standard scan speed in individual *en face* (xy axes) planes throughout the cellular z axis at 0.30 μm intervals.

### Transmission electron microscopy

The transmission electron microscopy was performed by harvesting 5×10⁵ mid-logarithmic phase trophozoites, and the cell pellet was fixed with 2.5% glutaraldehyde for 24 hours at 4°C as described previously [76, 77]. Post fixation, the pellet was resuspended in aqueous osmium tetroxide for 3 hours, followed by 6 washes with deionized distilled water. The cell pellet was then dehydrated in a series of alcohol, embedded in araldite resin, and incubated at 80°C for 72 hours for polymerization. An ultra-microtome (Leica Ultracut UCT-GA-D/E-1/00) was used to make 60 nm ultrathin sections, which were mounted onto nickel grids (G5526, Merck, USA); following this, blocking was carried out with PBSG (PBS with 0.2% gelatin and 0.5% BSA) for one hour at room temperature. The sections were then incubated with anti-RabD2 (1:30) primary antibody for 2 hours at room temperature, followed by 10 nm gold-conjugated secondary antibody (A31561 Invitrogen, USA) (1:50) for 1 hour at room temperature. The grids were incubated with saturated aqueous uranyl acetate and counterstained with Reynolds lead citrate to achieve contrast. The images were acquired using a Talos L120C transmission electron microscope (Thermo Fisher Scientific, USA) at CCMB, Hyderabad, India.

### Western blotting

*E. histolytica* trophozoites were harvested and lysed using the lysis buffer [50 mM Tris HCl, 150 mM NaCl, 1 mM DTT, 1 mM PMSF, 1% Triton X-100, 10 µM E-64 (E3132, Sigma-Aldrich, USA) and proteases inhibitor cocktail (04693116001, Sigma-Aldrich, USA)] at 4°C for 30 minutes. The whole cell lysate (25-35 µg) was resolved at 120V on a 12-15% acrylamide gel in reducing conditions. The proteins were electrophoretically transferred onto a PVDF membrane (10600023, Cytiva) at 90V for 90 minutes at 4°C. The transferred proteins on the membrane were incubated with blocking solution [5% (w/v) bovine albumin serum (BSAV-RO, Sigma-Aldrich, USA) in 1X PBS] for 1 hour at room temperature. The primary antibodies employed are anti-HA monoclonal (3724 Cell Signalling Technology) (1:1000), anti-RabD2 polyclonal (1:1000), anti-ubiquitin monoclonal (05-944, Merck, USA) (1:1000), and anti-CS1 polyclonal (1:1000) were incubated at 4°C overnight. Post incubation, the membrane was washed and incubated with horseradish peroxidase-conjugated anti-rabbit (111-035-144, Jackson Immuno Research Laboratories, USA) (1:1000) and anti-mouse (315-035-048, Jackson Immuno Research Laboratories, USA) (1:1000) followed by detection of immune-reactive proteins using chemi-luminescence (ImageQuant™ 500, USA).

### Treatment of sumoylation, ubiquitination inhibitors, reducing and chaotropic agents

To study the effect of sumoylation and ubiquitination inhibitors on amoebic parasites, 2x10^6^ trophozoites were incubated with 10 µM MG132 (M7449, Sigma-Aldrich, USA) or 50 µM 2-DO8 (SML1052, Sigma-Aldrich, USA) for 16 hours at 37 °C. The trophozoites were then lysed using lysis buffer followed by detection using western blotting as described above.

To study the RabD2 self-association, logarithmic phase wild-type trophozoites were harvested, lysed in lysis buffer and further incubated under rotation with a series of reducing and chaotropic agents; 1 mM or 10 mM DTT, 10% βME, 6M urea, 9N guanidine hydrochloride, respectively, for specified time points at 4°C. After each time point samples were prepared and subjected to western blotting.

### Adhesion Assay

The assay to investigate the adherence of *E. histolytica* trophozoites with CHO cells was adapted from a previous study [78]. Briefly, the amoebic trophozoites and CHO cells were harvested, washed, and co-incubated (1:20) in 1 ml PSF (calcium and magnesium free, Puck saline F) for 90 minutes on ice. Post incubation, the cells were centrifuged at 150x*g* for 5 minutes, and 0.8 ml of supernatant was discarded. The remaining cell pellet was gently broken down by rubbing the tube between hands and was transferred onto a clean 8-well slide. The cells were fixed with 4% PFA, and the slide was mounted with Prolong Diamond Antifade.

### Immunoblot quantification

The immunoblots were quantified using the ImageJ software by maintaining uniformity in the area and measuring the intensity, thereby deducing the background. The values obtained from ImageJ were utilised to plot the bar using GraphPad Prism 9, and their respective *p*-values are mentioned in the figure legend.

### RabD2 antibody production

The animal used in antibody generation study was in compliance with guidelines provided by the Committee for Control and Supervision of Experiments on Animals (CCSEA), Government of India and approved by the Institutional Animal Ethics Committee (protocol no. EAF/2023/PP/18). The full-length purified amoebic RabD2 protein was used as an antigen to produce the antibody. An eight-week old New Zealand rabbit was subcutaneously injected with a primary dose of 500 µg purified RabD2 protein prepared in Freund’s adjuvant (F5881, Sigma-Aldrich, USA). After 50 days, 750 µg of RabD2 protein was reinjected as the first booster dose. Later, a series of three extra booster doses with the same amount (750 µg) were injected with a time interval of 20 days each. Finally, serum was separated from the collected blood, and the antisera containing the anti-RabD2 antibody was further used for validation experiments. The pooled antisera containing the RabD2 antibodies were subjected to affinity purification using the protein A/G magnetic beads (88802, Thermo Fisher Scientific, USA) following the manufacturer’s instructions. The pre-immune serum was collected from the animal before injecting the RabD2 protein.

## Supporting information

Supporting Infromation

## Acknowledgments

This work was fully supported by grant number (EEQ/2019/000747) Science and Engineering Research Board, New Delhi, India to KV. We are sincerely grateful to Prof. Tomoyoshi Nozaki (Graduate School of Medicine, University of Tokyo, Japan), Prof. William A. Petri Jr. (University of Virginia, USA), and Prof. Sunando Datta (IISER, Bhopal, India) for kindly providing the anti-CS1, anti-Hgl, and anti-Rab5 antibodies, respectively. We also thank Shruti Shah (SERB-SSR trainee) for excellent technical help in protein purification.

