## Supporting Infromation for "Glutamic acid-lysine (EK) rich motif of RabD2 self-associates and regulates pathogenesis through multivesicular bodies pathway in *E. histolytica*": Supporting Information.docx

**S1 Fig.** **PCR amplification of amoebic RabD2 and sequence alignment with human Rab2A, Rab44, Rab45 and Rab46**

A. The putative amoebic RabD2 (EHI_164900) 795 bp PCR product was amplified using appropriate forward and reverse primers from cDNA pool of *E. histolytica*.

B. The cloned amoebic RabD2 DNA sequence was hypothetically translated into protein sequence and aligned with human Rab2A (P61019), Rab44 (Q7Z6P3), Rab45 (Q8IZ41) and Rab 46 (Q9BSW2). The amoebic RabD2 sequence contains unique 69 amino acids at the NH2-terminal and a conserved G domain (G1-G5 motif) in the COOH-terminal similar to that of the other Rab proteins.

**S2 Fig.** **Generation and validation of amoebic RabD2 antibody**

A. The coomassie gel image represents the purified Prescission Protease cleaved full length amoebic RabD2 protein (lane1 - 2.5 µg; lane2 - 5 µg) and GST-RabD2 protein (lane3 - 10 µg; lane4 - 20 µg). The purified protein shown in lane1 and lane2 was used in the immunisation of rabbit.

B. Immunoblotting was performed to validate the specificity of anti-RabD2 antibody. The pre-bleed (100 µg) sera (control) (collected from the rabbit prior to injecting the RabD2 protein) and *E. histolytica* whole cell lysate (100 µg) were resolved and probed with anti-RabD2 (1:1000) polyclonal sera. The *green* arrowhead marks the hypothetical molecular mass (approximately 29 kDa) and *red* arrowhead indicates higher molecular weight species (>30 kDa).

C. Wild type *E. histolytica* trophozoites were fixed, subjected to immunofluorescence and probed with pre-bleed sera (1:500) as control. Scale bar, 20µm.

D. Wild type *E. histolytica* trophozoites were probed with anti-RabD2 polyclonal sera (1:500, upper lane) and affinity purified sera (1:500, lower lane), followed by secondary antibody Alexa fluor-488. The confocal images show the fluorescence signals specific to endogenous RabD2 when probed with anti-RabD2 antibody. Scale bar, 20µm.

**S3 Fig. Evaluating the effect of reducing and chaotropic agents on endogenous RabD**2

A. Untransfected trophozoites lysate was prepared and incubated with denaturants; 1mM DTT, 10% β-ME, 6M urea and 9N guanidine hydrochloride for 30 minutes and subjected to western blotting using anti-RabD2 and anti-CS1 antibodies. The *green* arrowhead indicates the hypothetical molecular mass (approximately 29 kDa), the *blue* arrowhead indicates the highly intense mid-modified band (approximately 35 kDa) and the *red* arrowhead indicates higher molecular weight species (>37 kDa).

B. Untransfected trophozoites lysate was incubated for 16 hrs with 1 mM DTT, 10 mM DDT, 10% βME and 6M urea. The treated lysates were subjected to western blot and probed with anti-RabD2 and anti-CS1 antibodies. The *green* arrowhead indicates the hypothetical molecular mass (approximately 29 kDa), the *blue* arrow head indicates the highly intense mid-modified band (approximately 35 kDa) and the *red* arrowhead indicates higher molecular weight species (>37 kDa).

**S4 Fig. Localization of amoebic RabD2 and amoebic Rab5 on MVB-like structures**

A. Association of endogenous RabD2 with MVB-like structures. Wild type trophozoites were subjected to immunofluorescence assay using anti-RabD2 antibody and imaged using structured illumination microscope (SIM). The square box represents the zoomed panel for the localization of RabD2 on MVBs like structures. The *white* arrowheads show the presence of RabD2 on the surface of MVBs and the *white* arrows indicate RabD2 packed within the MVB. Scale bar, 20µm.

B. Wild type HA-RabD2 associates with endogenous Rab5 on MVBs. The wild type HA-RabD2 trophozoites were probed with anti-HA and anti-Rab5 antibodies. The square box represents the zoomed panel which shows that RabD2 is colocalized with Rab5 on the surface (*white* arrowheads) and within the MVB like structure (*white* arrow). The localisation of HA-RabD2 on the MVB surface (*green* arrowhead) and ILV (*green* arrow) accompanied with Rab5 localisation on the MVB surface (*red* arrow heads) and its ILV (*red* arrow) are depicted in this image. Scale bar, 20µm.

**S5 Fig.** **HA-RabD2 overexpression induces the MVBs and ubiquitination**

A. pEhTex-HA and HA-RabD2 expressing cells were subjected to transmission electron microscopy. Scale bar, 5 µm.

B. The number of MVBs from the TEM images were manually quantified and the graph plotted represents the mean ± S.D. Statistical significance was calculated using unpaired two-tailed Student’s t test (n=10 cells/construct).

C. To Evaluate the levels of ubiquitination upon the wild type HA-RabD2 overexpression. The pEhTex-HA and HA-RabD2 (in absence and presence of tetracycline) expressing trophozoites were lysed and subjected to immunoblotting with anti-ubiquitin, anti-HA and anti-CS1 antibodies. An increase in the levels of global ubiquitination was observed in tetracycline induced HA-RabD2.

**S6 Fig.** **Effect of RabD2 and its mutant trophozoites on the adherence with the CHO cells**

Wild type HA-RabD2 and its mutants HA-RabD2QL, HA-RabD2TN and HA-Δ69RabD2 were subjected to adhesion assay by incubating the trophozoites with CHO cell for 90 minutes. The cells were fixed and images were captured. Eh: *E. histolytica,* CHO: Chinese hamster ovary. Scale bar, 20µm.

**Supporting methodology**

**Cloning expression and purification of amoebic RabD2 protein from *E. coli***

The pGEX-6p (28954648, Cytiva, USA) plasmid was digested with *BamHI* and *Xhol*. Similarly, full-length RabD2 was re-amplified from the confirmed clone (pEhTex-HA-RabD2) using the primers with restriction sites *BamHI* and *XhoI*. The digested HA-RabD2 and pGEX-6p were ligated using T4 DNA ligase (M0202S, New England Biolabs, USA). The ligated product was transformed to *E. coli* DH5-α and recombinant clones (pGEX-6p-RabD2) were confirmed by unidirectional sanger sequencing. The recombinant expression plasmid pGEX-6p-RabD2 was transformed into *E. coli* BL-21 DE3. Briefly, 600ml of the bacterial culture was grown at 37˚C until the optical density (600 nm) reached 0.4, then the expression of fusion protein was induced by adding 0.25 mM Isopropyl β-D-1-thiogalactopyranoside (I6758, Sigma-Aldrich, USA) and incubation was carried over for 4 hours. The cells were pelleted down by centrifugation at 5000 rpm for 10 minutes at room temperature. The supernatant was discarded and the pellet was resuspended in 10 ml lysis buffer [(100 mM NaCl, 2 mM EDTA, 20 mM HEPES pH 6.8, 5 mM DTT) supplemented with cOmplete™ ULTRA protease inhibitor cocktail (05892970001, Roche, USA). The re-suspended cells were sonicated at a frequency of 20kHz for 30 minutes and the whole lysate was centrifuged at 16000x*g* for 20 minutes at 4˚C. The supernatant was collected and supplemented with 1% Triton X-100. Later, 500 µl of glutathione sepharose 4B beads (GE17075601, Cytiva, USA) were washed thrice with the above prepared lysis buffer by centrifuging at 500x*g* for 5 minutes at 4˚C. Finally, washed beads were resuspend in lysis buffer supplemented with 1% Triton X-100. The glutathione sepharose beads were added to the above supernatant (containing the recombinant protein) and placed on an end over end rotamer for overnight binding at 4˚C. After binding, the remaining supernatant was removed by centrifugation at 5000 rpm for 10 minutes and the beads (binding with the recombinant protein) were incubated with elution buffer (50 mM Tris, 150 mM NaCl, 1 mM EDTA, 1 mM DTT and 20% glycerol) containing precision protease (20 µl PreScission Protease in 500 µl buffer) (GE27084301, Cytiva, USA) at 4˚C for overnight. After protease treatment the beads were centrifuged at 5000 rpm for 10 minutes at 4˚C and the supernatant containing the purified RabD2 protein was collected into a clean 1.5ml microcentrifuge tube. The purity of RabD2 protein was evaluated by SDS-PAGE analysis and quantified using BSA standards (23209, Thermo Fisher Scientific, USA) and bradford protein assay (23236, Thermo Fisher Scientific, USA).
