## Supplementary figures and images for "Glutamic acid-lysine (EK) rich motif of RabD2 self-associates and regulates pathogenesis through multivesicular bodies pathway in *E. histolytica*"

### S1_Fig.tif

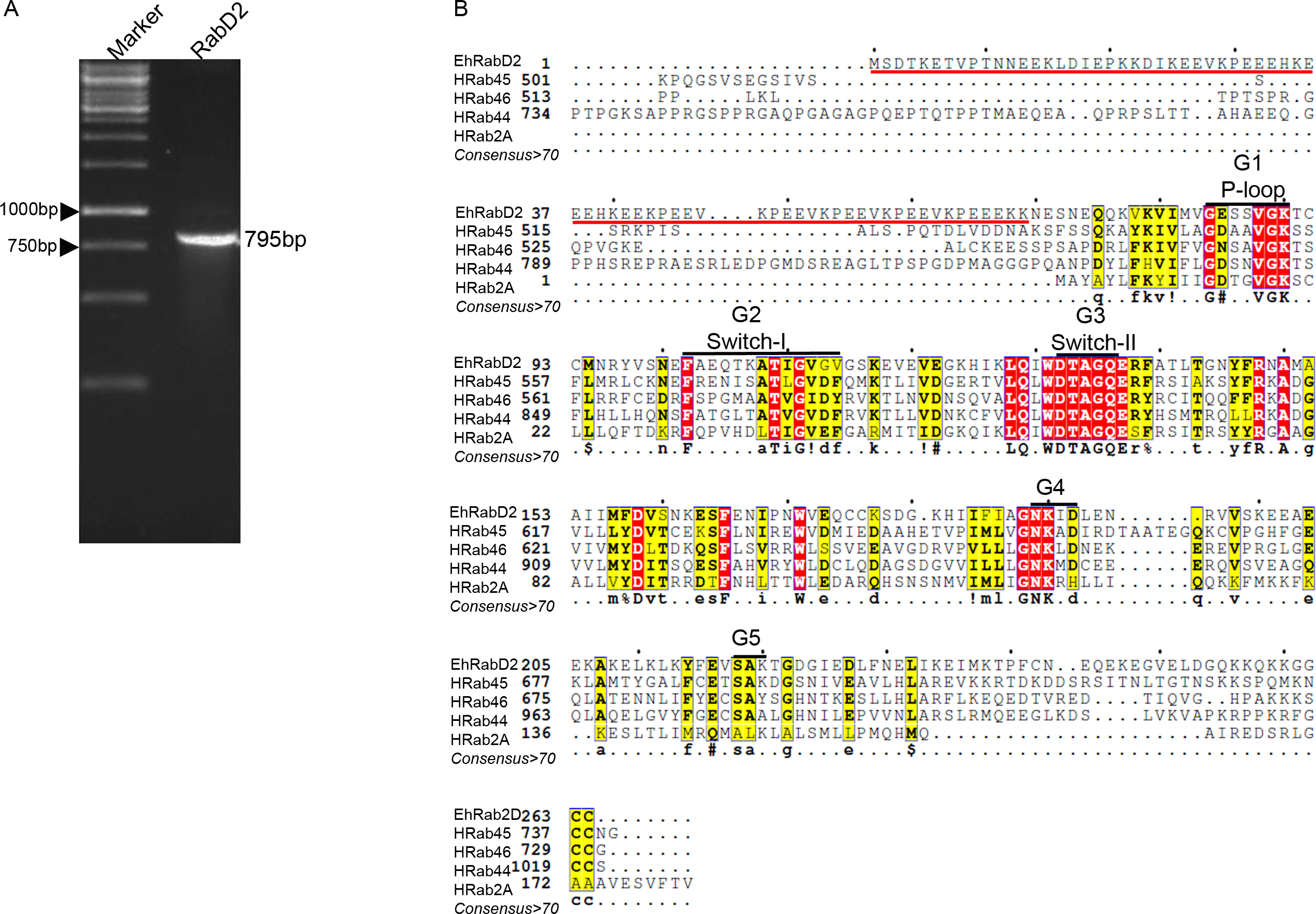

### S2_Fig.tif

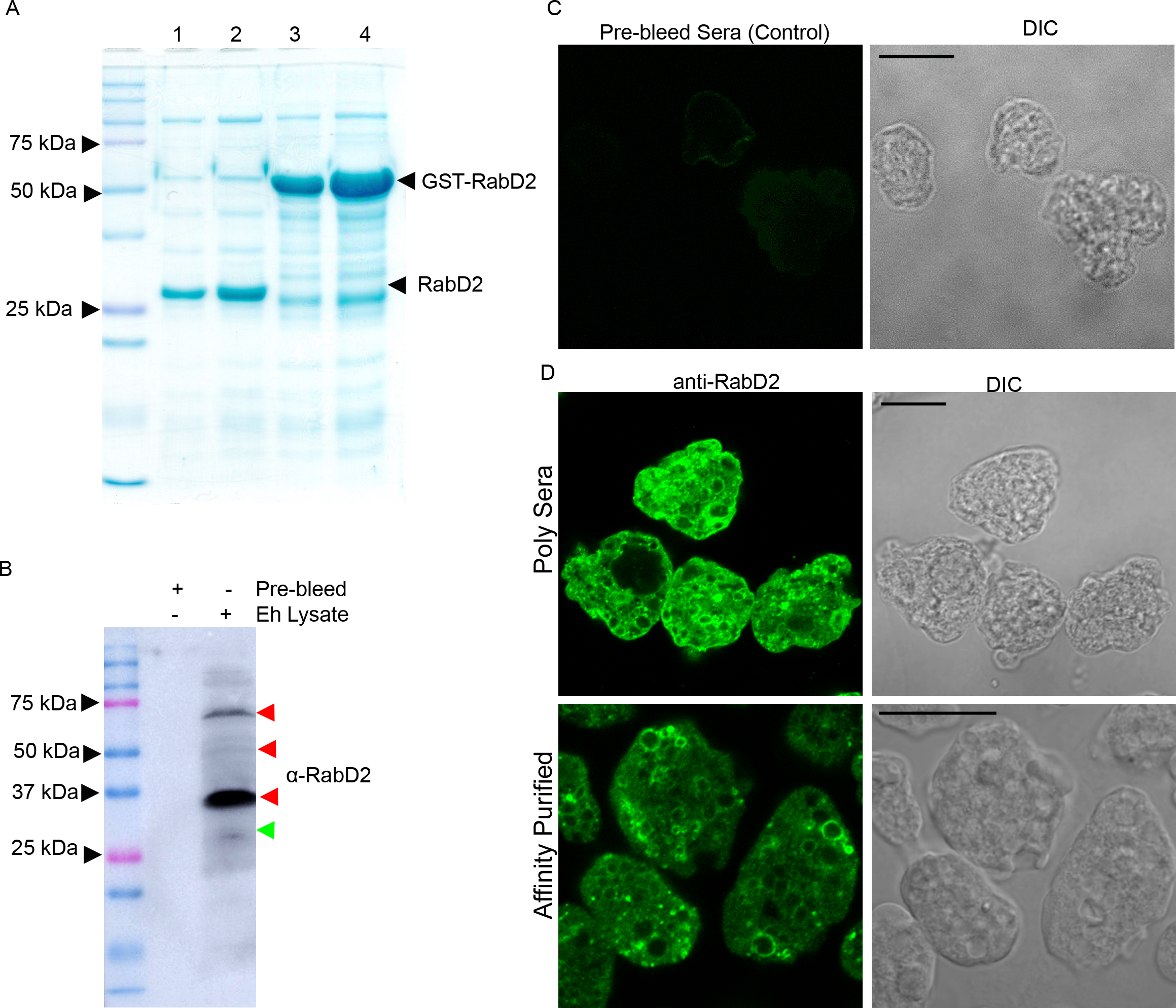

### S3_Fig.tif

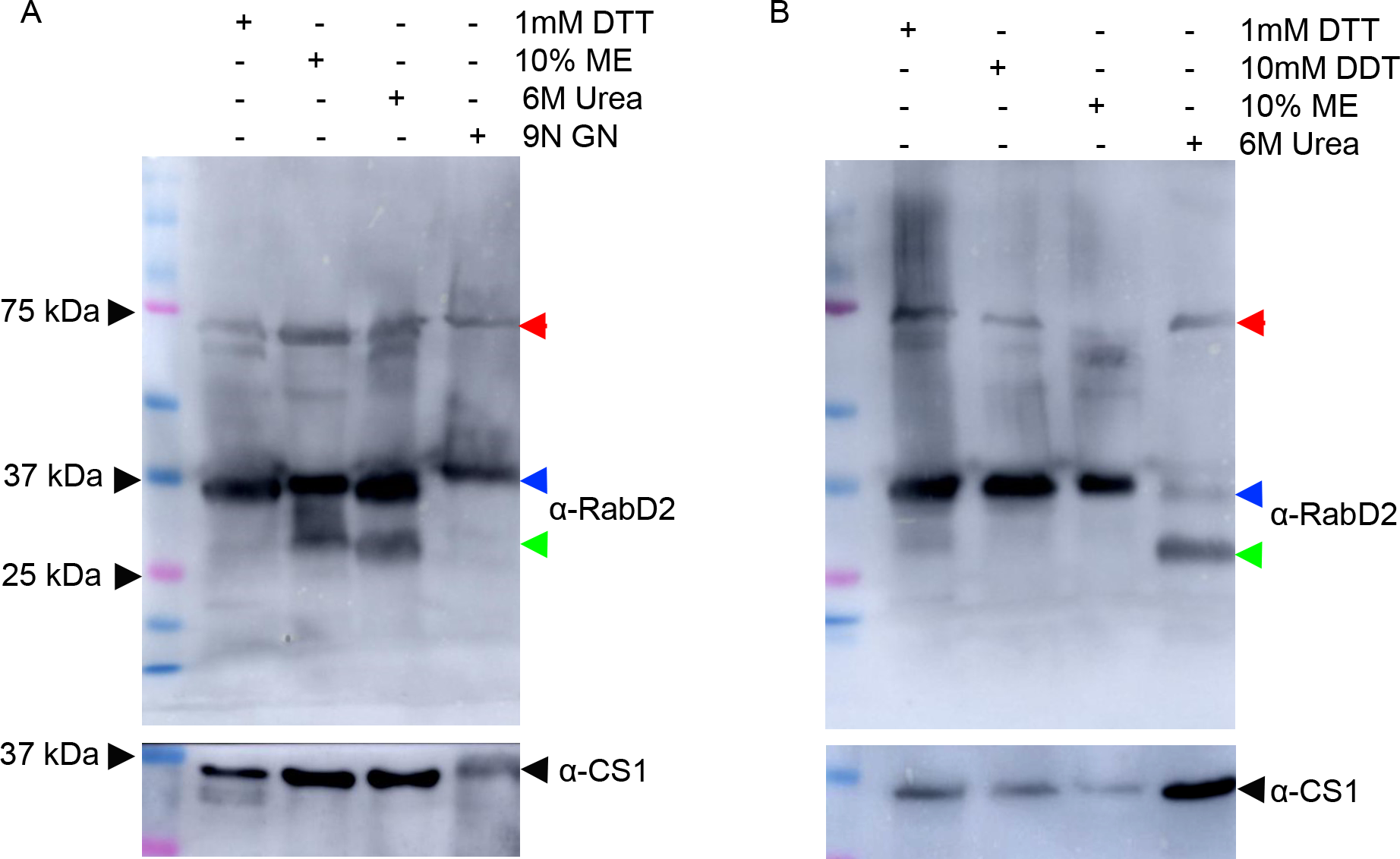

### S4_Fig.tif

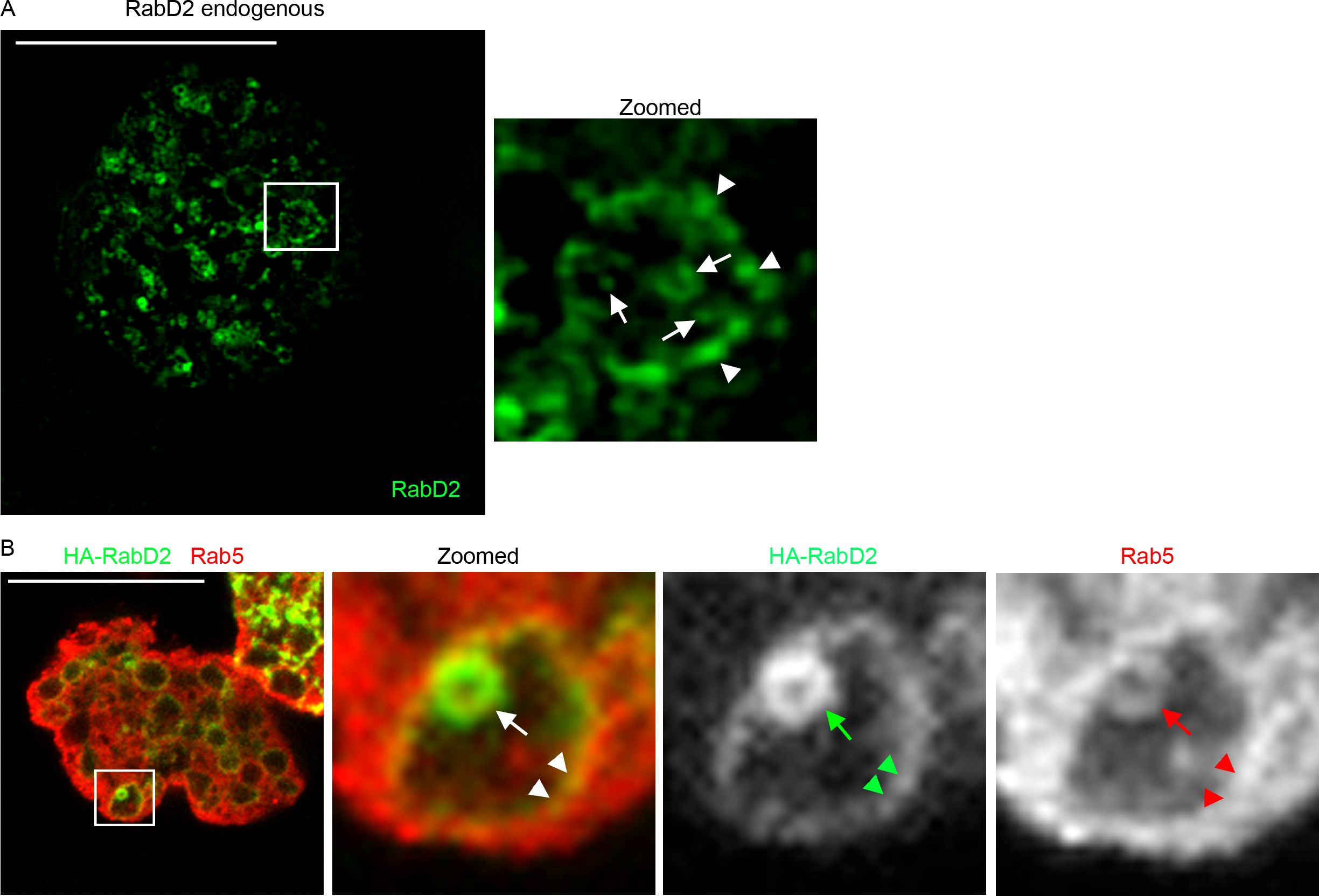

### S5_Fig.tif

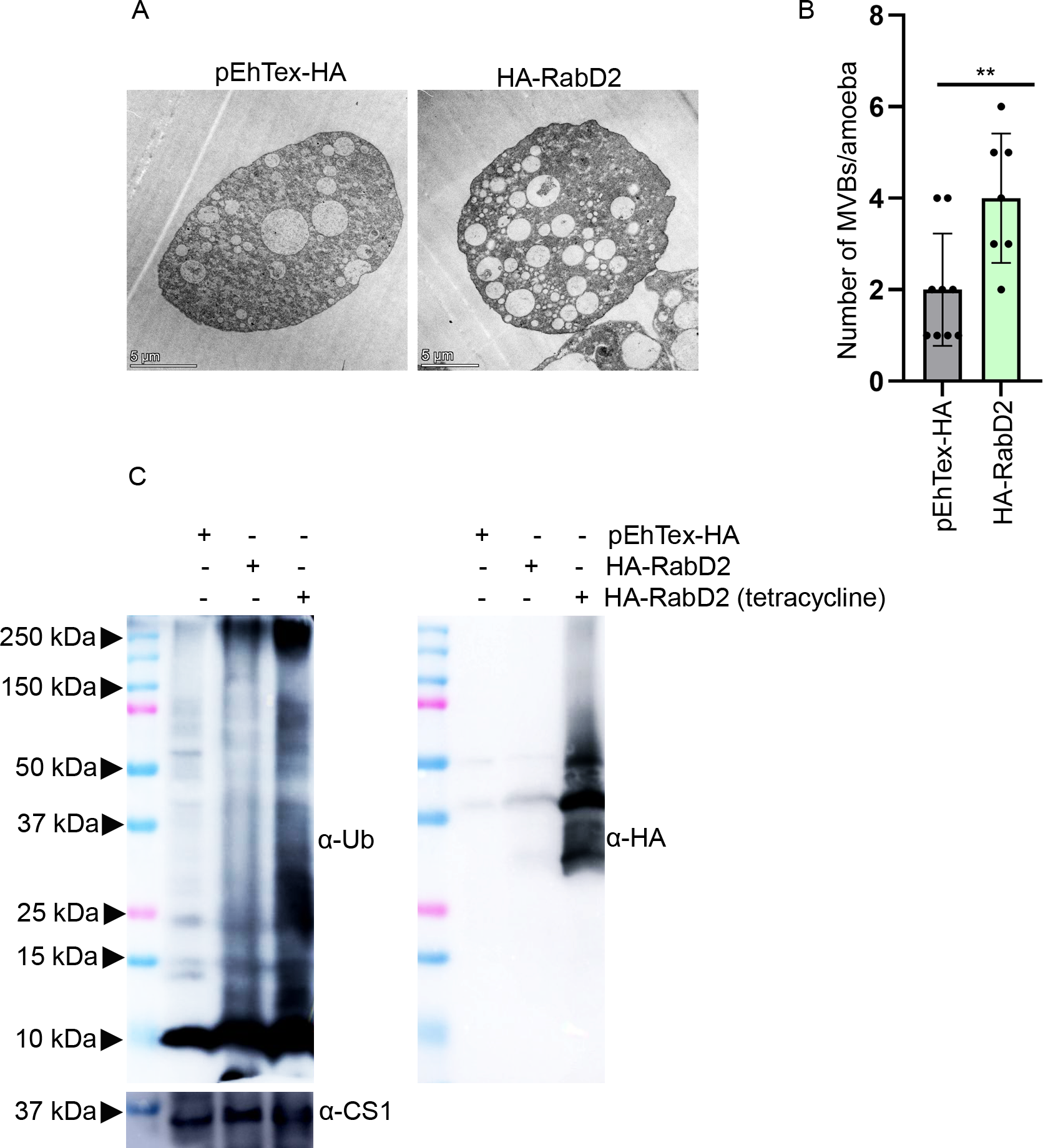

### S6_Fig.tif

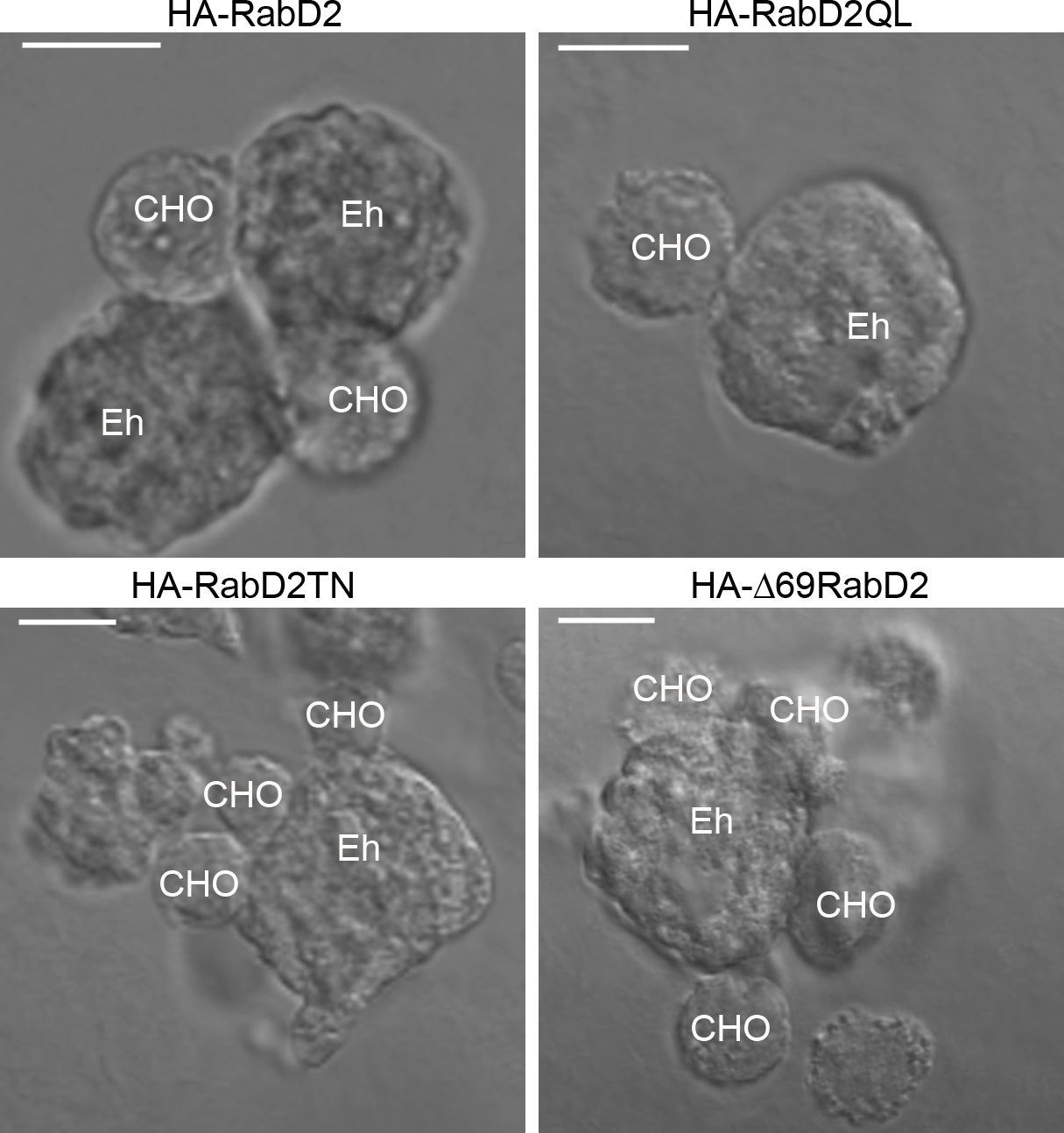
